# Adipose Tissue Overexpression of Nicotinamide Phosphoribosyltransferase Prevents Metabolic Dysfunction in Obese Mice

**DOI:** 10.1101/2025.09.30.679563

**Authors:** Daniel Ferguson, Elise I. Gadson, Kathleen R. Markan, Jun Yoshino, Maxwell Lin, Mohammad Habibi, Snigdha Tiash, Xiaoxia Cui, Evguenia Kouranova, Edziu Franczak, Qiuyuan Guo, Jill Kealing, Terri A. Pietka, Kim H. H. Liss, John P. Thyfault, Gary J. Patti, Brian N. Finck, Clair Crewe, Sandip Mukherjee

**Author notes:** Corresponding author: Sandip Mukherjee.

## Abstract

Nicotinamide adenine dinucleotide (NAD+) is a vital coenzyme and a central factor in energy metabolism. Nicotinamide phosphoribosyltransferase (NAMPT) maintains the cellular NAD+ pool by synthesizing the NAD+ precursor, nicotinamide mononucleotide (NMN), and diminished adipocyte NAMPT activity has been implicated in aging- and obesity-related metabolic dysfunction. Herein, we examined the effects of overexpressing or knocking out NAMPT in adipocytes on metabolic dysfunction and interorgan communication in mice. We generated new adipocyte-specific NAMPT overexpressing(*ANOV*) mice model. Male *ANOV* mice are protected from diet-induced metabolic dysfunction including adipose tissue inflammation, glucose intolerance, and insulin resistance. In contrast female *ANOV* mice were less protected from metabolic dysfunction, possibly due to higher endogenous expression of NAMPT in *WT* female mice. Livers of *ANOV* mice showed improved insulin signaling, increased NAD content, and reduced steatosis, suggesting that NAMPT regulates interorgan communication between adipocytes and hepatocytes. Extracellular vesicles (EV) isolated from *ANOV* mice enhanced insulin signaling in HepG2 cells and improved glucose tolerance in *WT* obese mice. In contrast, EV from *ANKO* mice suppressed HepG2 insulin signaling and inhibition of EV release improved glucose tolerance in *ANKO* female mice. Collectively, these data highlight a novel mechanism by which adipocyte NAD+ metabolism regulates systemic metabolic dysfunction via EVs.

## INTRODUCTION

Nicotinamide adenine dinucleotide (NAD+) is an enzyme cofactor required for many metabolic reactions critical for maintaining cellular energy homeostasis. For example, reducing NAD+ to NADH transfers chemical energy stored in nutrients to the electron transport chain to fuel ATP synthesis. NAD+ also regulates the activity of enzymes that control various signaling pathways such as the sirtuin family of deacetylases. NAD+ abundance is diminished in obesity-related metabolic diseases and aging while supplementation of NAD+ precursors improves metabolic function, insulin sensitivity, and may have anti-aging effects[1–6].

A high ratio of NAD+ to NADH energetically favors the reduction of NAD+ to NADH. The cell maintains this redox state by several mechanisms, including the activity of NAD+ generating reactions such as the conversion of pyruvate to lactate by lactate dehydrogenase [6, 7]. Additionally, NAD+ can be synthesized *de novo* from amino acids or produced by a salvage pathway from nicotinamide riboside (NR) and nicotinamide mononucleotide (NMN) [8]. Nicotinamide phosphoribosyltransferase (NAMPT) catalyzes a key step in the salvage pathway by converting nicotinamide to NMN [9–12]. NAMPT expression is reduced in obesity and aging, coincident with diminished NAD+ abundance[1, 2, 13]. Mice lacking NAMPT in adipocytes (*ANKO* mice) exhibit insulin resistance and metabolic dysfunction, and NMN supplementation reverses the effects of NAMPT deficiency[4, 14, 15].

NAD+ insufficiency results in impaired mitochondrial function and dysregulated metabolism [2, 4, 14, 16, 17]. NAMPT deficiency also leads to impairments in sirtuin activity and suppression of PPAR signaling in adipocytes [14]. Accordingly, therapeutic approaches to enhance sirtuin or PPAR activity can also reverse metabolic dysfunction in *ANKO* mice. Furthermore, NAMPT circulates in extracellular vesicles (EV) released by adipocytes and treating aging mice with NAMPT-containing EV delays aging and extends lifespan[17–19]. However, the effects of circulating NAMPT-containing EV on the development of metabolic dysfunction are poorly understood.

In this work, we examined the effects of overexpressing NAMPT in adipocytes in mice (*ANOV* mice) and identified novel mechanisms by which adipocyte NAMPT activity regulates systemic metabolic homeostasis and liver metabolism.

## RESEARCH DESIGN AND METHODS

Additional methodological details and procedures can be found in Supplemental Materials.

### Animal studies

All animal studies were approved by the Institutional Animal Care and Use Committee of Washington University in St. Louis. All methods are reported in accordance with ARRIVE guidelines for the reporting of animal experiments. Mice were housed at 23°C under 12-hour light/dark cycles and given *ad libitum* access to food and water throughout the duration of treatment. For diet studies, starting at 8 weeks of age, male mice were fed high-fat diet (HFD; 60% kcal from fat, Research Diets, D12492) or a matched low-fat diet (LFD; 10% kcal from fat, Research Diets, D12450J) for 12 weeks. Due to the sexual dimorphic response to HFD[20, 21], female mice were fed LFD or HFD for 16 weeks. Mice were fasted for 4-5 h prior to euthanasia by carbon dioxide asphyxiation. Plasma and tissues were collected, weighed, and preserved following standard procedures for the assessment of various parameters.

For a separate study, after twelve weeks on HFD (60% kcal from fat, Research Diets, D12492) C57BL6/J mice were injected intravenously once a week for 3 weeks with 1 × 10^9^ EVs isolated from adipose tissue of chow-fed male *WT* or *ANOV* mice[22, 23]. Mice then underwent glucose tolerance testing (GTT) and were euthanized for blood and tissue collection.

Sixteen-week-old chow-fed female *WT* or *ANKO* mice were injected intraperitoneally with GW4869 (60 mg/mouse) or vehicle twice a week for 6 weeks. Mice then underwent glucose tolerance testing (GTT) and were euthanized for blood and tissue collection.

### EV treatment

For in vitro cell culture experiments, hepatocytes (HepG2) were treated with *WT* or *ANOV* mice adipose tissue-derived EV (1x 10^8^ / well) for 48 hours followed serum-starvation for 4 h and insulin (100 nM) treatment for 5 min. Number of EVs were counted prior to treatment using NanoSight analysis and dose of EV treatment was based on an earlier publication[24].

### Plasma EV characterization by flow cytometry

For total EV quantification, 2 µl plasma was combined with 25 µM carboxyfluorescein succinimidyl ester (CFSE) diluted in PBS. CFSE stain was used to label and track EVs in plasma following previous protocol[25]. Briefly, samples were incubated for 30 minutes at 37°C. EVs were purified from free dye and plasma proteins with size exclusion chromatography, as described for ex vivo-derived EV. For detection of NAMPT in plasma EVs, 2 µl plasma was mixed with FcBlock (1:200) and 0.00001% triton X-100 in a 25 µl volume for 10 minutes. Anti-NAMPT antibody (Cat # A300-372A; Bethyl Laboratories) or rabbit Isotype control antibody conjugated to CF633 (Biotium) was added to samples for a final dilution of 1:100. Samples were incubated rocking overnight at 4°C. Flow cytometry analysis was conducted with the Cytek NorthernLights spectral flow cytometer with enhanced small particle detection (ESP). Purified EV from mouse plasma were titrated to determine the appropriate dilution to prevent swarming (abort rate remained under 10% of the event rate). The violet side scatter laser was used as the trigger channel and the threshold was set to 600. The side scatter trigger channel was calibrated for EV size determination with Rosetta Calibration beads (Exometry) using the recommended parameters (EV model/refractive index). NAMPT containing EVs were gated based on rabbit Isotype control antibody for further downstream analysis (Supplemental Fig. 6B).

### Statistics

All data are presented as mean ± SEM. Significance was determined by two-tailed Student t test or in the case of systemic assays, two-way ANOVA. P < 0.05 was considered statistically significant. All quantitative PCR data points were the average of technical duplicates. For all mouse studies, the n value corresponds to individual mice of a given treatment. Data were analyzed using GraphPad Prism software.

## RESULTS

### Generation of adipocyte NAMPT overexpressing (ANOV) mice

To assess the effects of adipocyte-specific NAMPT overexpression, we generated mice with the human NAMPT coding sequence inserted at the C-terminus of *Adipoq*, resulting in NAMPT expression driven by the endogenous adiponectin promoter (Supplemental Fig. 1a). Accordingly, there was an increase in NAMPT protein abundance by more than 3-fold in epididymal white adipose tissue (eWAT) and inguinal white adipose tissue (iWAT) of male *ANOV* transgenic mice compared to wild-type (*WT*) littermate control mice (Fig.1a-b and Supplemental Fig. 1b-c). Unexpectedly, NAMPT was not overexpressed in brown adipose tissue (BAT) in *ANOV* mice (Supplemental Fig. 1c), which may be partially explained by the observation that NAMPT expression is markedly higher in BAT than WAT in *WT* mice (Supplemental Figure 1c). NAMPT was not overexpressed in other tissues including liver, muscle, and kidney of *ANOV* mice (Fig. 1b). On a chow diet, male and female *ANOV* mice were overtly normal and exhibited no changes in body weight or composition (Fig.1c). Compared to *WT* littermate controls, 16 h fasting blood glucose concentrations were lower in 2-month-old male *ANOV* mice, but not in female *ANOV* mice, compared to sex-matched littermate *WT* mice (Fig.1d). In 10-month-old male mice maintained on a chow diet, glucose tolerance was improved in *ANOV* mice compared to *WT* littermates (Fig.1e), despite no differences in body composition (Supplemental Fig.1d).

**Fig. 1.**
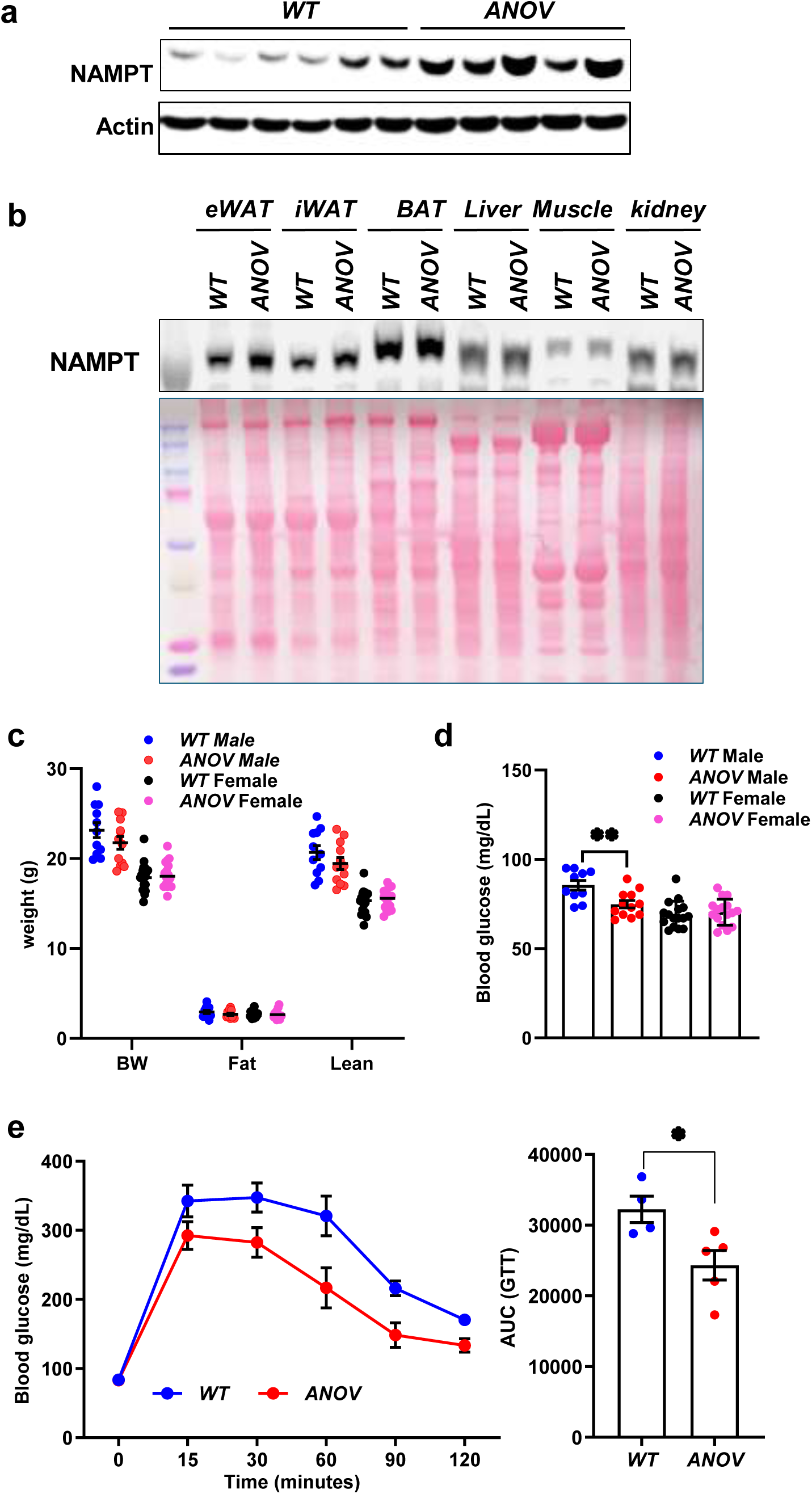
Generation of adipocyte NAMPT overexpressing (*ANOV*) mice. **(a)** Immunoblot showing protein levels of NAMPT in epididymal white adipose tissue (eWAT) of chow-fed, *WT* and *ANOV* mice (n=5-6). Actin was used as loading control. **(b)** Protein level of NAMPT in the indicated tissues of *WT* and *ANOV* mice. Ponceau was used for total protein content. **(c).** Body composition was determined by ECHO MRI in chow-fed male and female mice (N=14-19 / cohort). **(d)** Blood glucose levels were measured in chow-fed male and female mice after 16 h fasting. **(e)** Chow fed 10-months old male mice were subjected to glucose tolerance testing (GTT) (n=4-5). Area under the curve (AUC) was calculated based on GTT (n=4-5). Data represented as mean ± SEM. Significance determined by Student’s t-test. *P < 0.05, **P < 0.01.

### ANOV mice are protected from high-fat diet-induced metabolic dysfunction

To determine how adipocyte-specific NAMPT overexpression would affect diet-induced metabolic dysfunction, we placed littermate male *WT* and *ANOV* mice on a high-fat diet (HFD; 60% kcal from fat) or low-fat diet (LFD; 10% kcal from fat) for 12 weeks. No difference in body weight or composition was noted between *ANOV* or *WT* mice on either the LFD or HFD (Fig. 2a and Supplemental Fig. 2a). On the LFD, there was a modest increase in plasma adiponectin levels in *ANOV* mice compared to *WT*, but plasma lipids were unchanged (Supplemental Fig. 2b-d). On HFD, *ANOV* mice had a strong trend towards decreased plasma triglycerides compared to *WT* mice, but there were no differences in circulating adiponectin levels (Supplemental Fig. 2b-c). Fasting plasma insulin (Fig.2b) and glucose (Supplemental Fig.2e) concentrations were significantly lower in HFD fed *ANOV* mice relative to *WT* mice. Male *ANOV* mice also had improved glucose tolerance (Fig. 2c) on both diets, and insulin tolerance (Fig. 2d) on HFD. Collectively, these data suggest that male *ANOV* mice are protected from HFD-induced glucose and insulin intolerance.

**Fig. 2.**
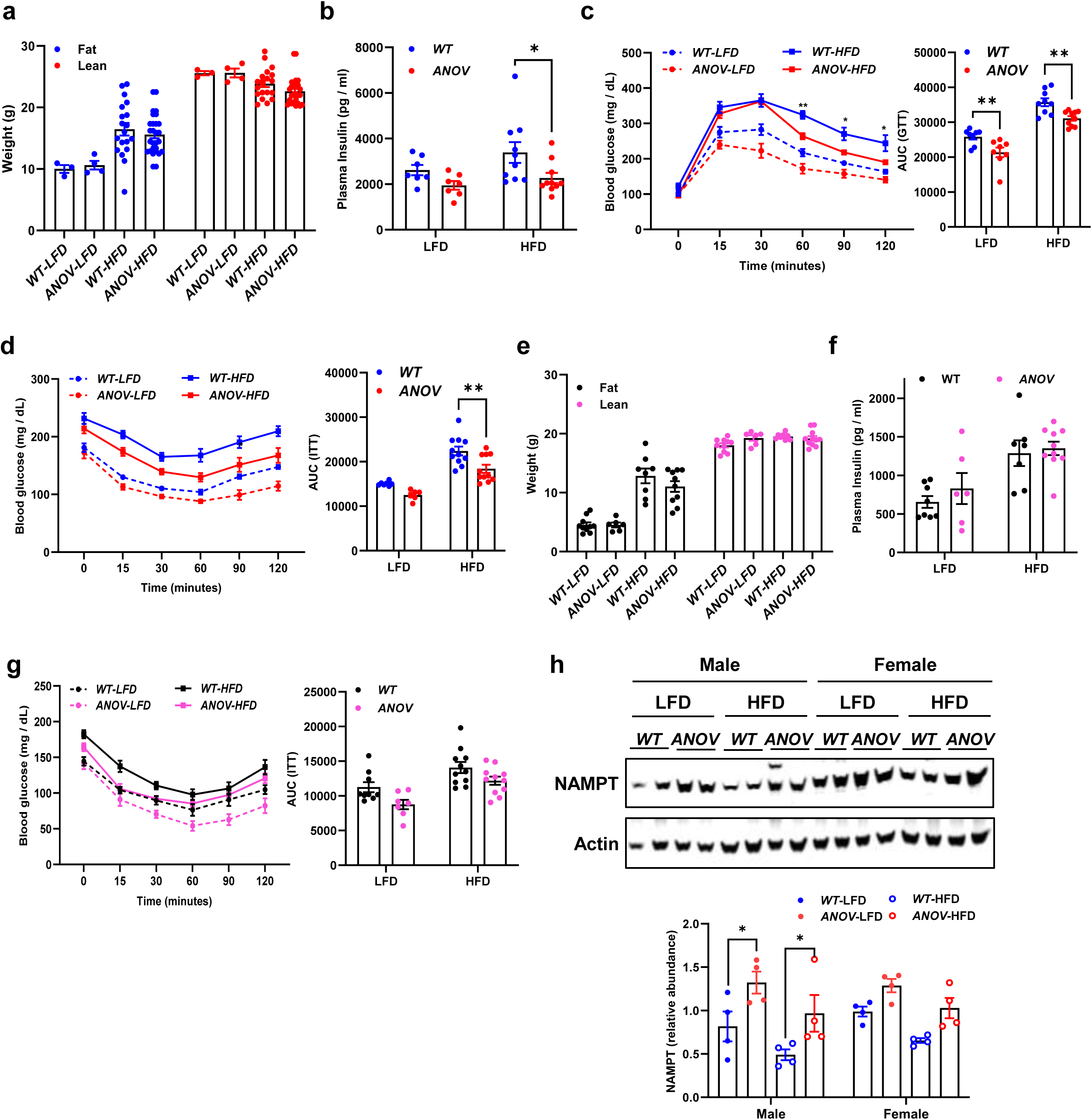
*ANOV* mice are protected from high-fat diet-induced metabolic dysfunction. *WT* and *ANOV* mice were fed with either a LFD and HFD for 12 (male) or 16 (female) weeks, followed by subsequent analyses. **(a)** Body composition in diet-fed male mice was determined by ECHO MRI (LFD n=3-4; HFD n=19-28). **(b)** Plasma insulin levels were determined in male mice following a 4 h fast (LFD n=6-7; HFD n= 10). **(c)** GTT was performed after 8 weeks of diet feeding in male mice (LFD n=8; HFD n=8-9). **(d)** ITT was performed after 10 weeks of diet feeding in male mice (LFD n=7-9; HFD n=11). **(e)** Body composition of LFD and HFD-fed female mice was determined by ECHO MRI. (n=8-11).**(f)**. Plasma insulin concentrations were determined in 4-h fasted female mice (LFD n=6-8; HFD n= 7-10).**(g)** ITT was performed after 14 weeks of diet in female mice (LFD n=7-9; HFD n=11). **(h)** Immunoblot showing protein level of NAMPT in white adipose tissue (WAT) of LFD or HFD *WT* and *ANOV* male and female mice (n=2). Actin was used as loading control. The number of mice (n) is presented as individual datapoints. Mean ± s.e.m. shown within dot plots. For multiple comparisons, two-way ANOVA with Holm-Sidák multiple comparison test was performed. *P < 0.05, **P < 0.01.

In female mice on HFD for 16 weeks, NAMPT genotype did not affect body weight, body composition, or plasma insulin concentrations (Fig. 2e-f and Supplemental Fig.2f). However, there was a trend towards improved insulin tolerance in female *ANOV* mice compared to *WT* mice on either diet (Fig.2g). Interestingly, we observed that *WT* female mice tended to have higher expression of NAMPT in gonadal WAT compared to *WT* males, suggesting an effect of sex on the regulation of NAMPT protein expression. NAMPT protein abundance was not significantly increased in *ANOV* female WAT compared to *WT* female mice on either diet (Fig. 2h), which may be explained by the higher endogenous expression of NAMPT.

### Adipose tissue NAMPT abundance is regulated by estrogen

To investigate the effects of estrogen on NAMPT expression, sham or ovariectomized (*OVX*) C57BL6/J female mice were placed on HFD for 4 weeks. Ovariectomized mice exhibited significantly less NAMPT protein expression in gonadal WAT (Fig. 3a). In a separate experiment, ovariectomized C57BL6/J mice on HFD were administered estradiol or vehicle (sesame oil). Interestingly, estradiol supplementation for 4 weeks significantly increased NAMPT expression compared to vehicle controls (Fig. 3b). These results suggest that adipose tissue NAMPT protein abundance is regulated by estrogen levels.

**Fig. 3.**
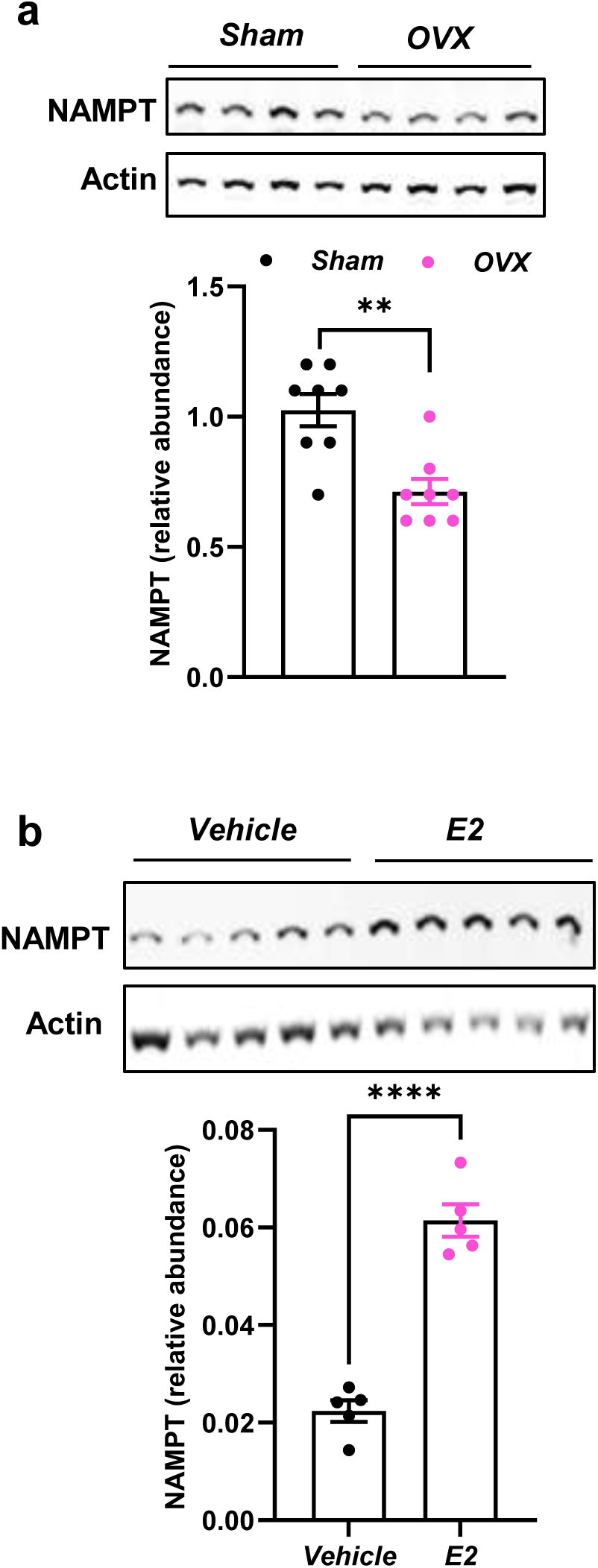
Adipose tissue NAMPT abundance is regulated by estrogen. Immunoblot showing protein abundance of NAMPT in gonadal fat tissue of HFD fed **(a)** *Sham* and *ovariectomized (OVX)* female mice (n=8) or **(b)** vehicle or estradiol supplemented (E2) OVX mice (n=5). Actin was used as loading control.

### Adipose tissue inflammation is reduced in ANOV mice on a high-fat diet

Next, we performed bulk RNA sequencing (RNAseq) on epididymal adipose tissue from male *WT* and *ANOV* mice fed either the LFD or HFD. On LFD, the expression of only 15 genes was significantly changed (13 decreased and 2 increased) in *ANOV* mice compared to *WT* mice (Supplemental Fig. 3a-b). However, on HFD, 91 genes were significantly different (75 decreased and 16 increased) in *ANOV* mice compared to *WT* mice (Fig. 4a and b). Pathway analysis of Hallmark gene sets identified adipogenesis and oxidative phosphorylation as being upregulated by NAMPT overexpression in both the LFD (Supplementary Fig. 3c) and HFD groups (Fig. 4c). Consistent with this, qPCR data demonstrated that *Pparg, Slc2a4* (GLUT4), and *Sirt1* expression was increased in *ANOV* mice on HFD, while *Ppargc1a* expression tended to increase (Fig. 4d). Furthermore, several genes associated with insulin sensitivity were upregulated (*Lpin1, Fam13A, Rbp7*) [26–28] whereas inflammatory genes involved in metabolic dysfunction (*Il1rn* and *Ccl7*) [29–31] were downregulated in HFD fed *ANOV* mice adipose tissue (Fig. 4e). Interestingly, one of the top upregulated genes in *ANOV* mice adipose tissue was *Lpin1*, which we have recently shown to be important for adipocyte function and systemic metabolism in both mice and humans[26, 32].

**Fig. 4.**
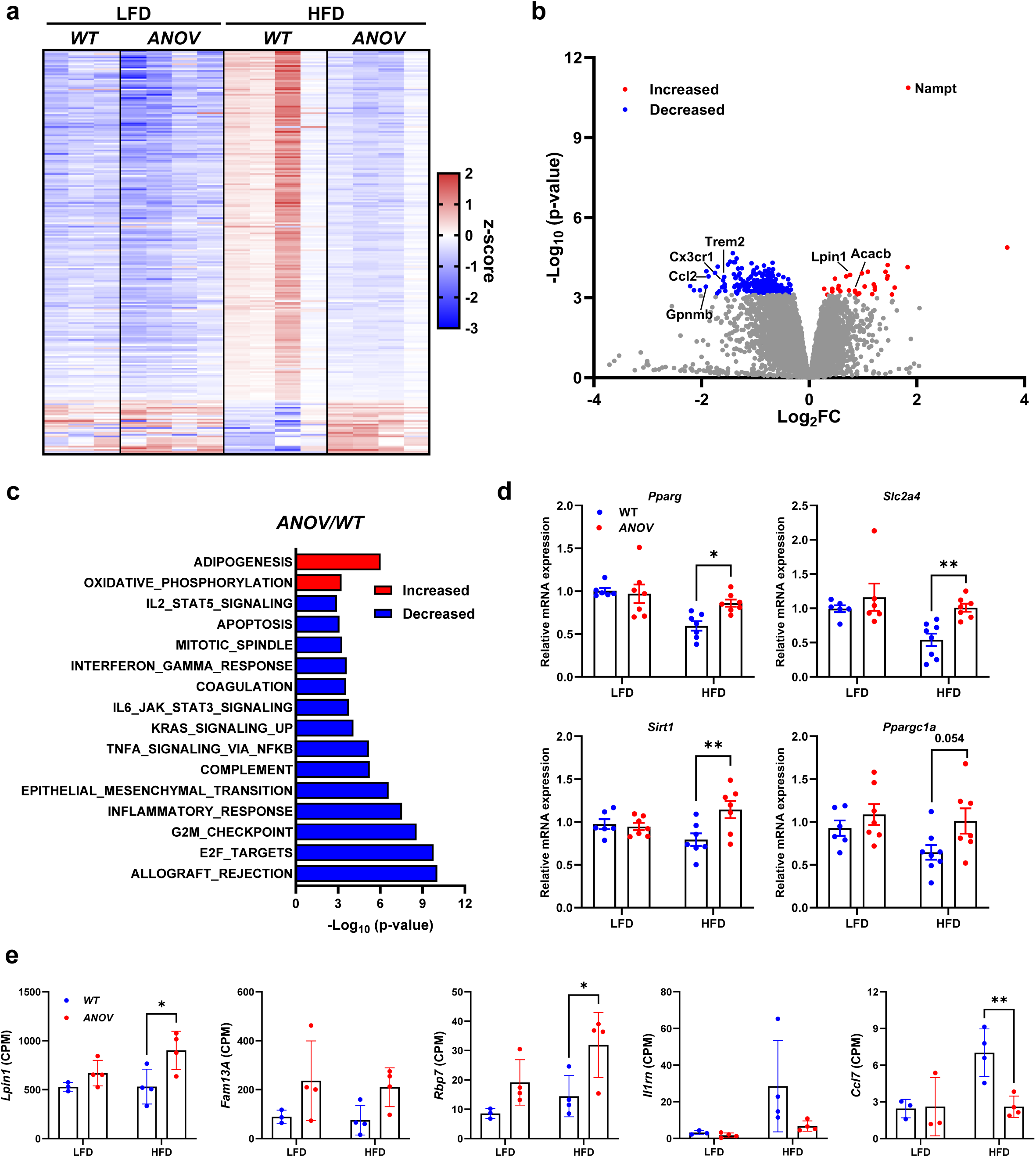
RNA sequencing reveals suppression of inflammatory pathways in *ANOV* adipose tissue on a high fat diet. Bulk RNA sequencing was performed on epididymal white adipose tissue from *WT* and *ANOV* male mice after 12 weeks of either LFD or HFD. **(a).** Heatmap representation of gene expression across all groups of genes that were significantly differentially expressed (adj. P value < 0.05) in HFD-fed *ANOV* vs HFD-fed *WT*. **(b).** Volcano plot of differentially expressed genes in HFD-fed *ANOV* vs HFD-fed *WT* with genes that were significantly downregulated or upregulated highlighted in blue or red, respectively (adj. P value < 0.05). **(c).** Hallmark gene sets that were significantly different (false discovery rate < 0.05) and substantially increased (red; LogFC > 3) or decreased (blue; LogFC < −3) when comparing HFD-fed *ANOV* vs HFD-fed *WT*. **(d).** Relative mRNA expression of genes associated with adipocyte differentiation (*Pparg*), glucose metabolism (*Slc2a4*), and mitochondrial metabolism (*Ppargc1a* and *Sirt1*) in eWAT of *WT* and *ANOV* mice by quantitative RT-PCR (LFD n=6-7; HFD n=7-8). **(e)**. Data from individual genes identified by bulk RNA sequencing. Data represented as counts per million (CPM). The number of mice (n) is presented as individual datapoints. Mean ± s.e.m. shown within dot plots. For multiple comparisons, two-way ANOVA with Holm-Sidák multiple comparison test was performed. *P < 0.05, **P < 0.01.

Pathway analysis also revealed the reduction of several inflammatory and immune responses in *ANOV* mice compared to *WT* mice on HFD (Fig. 4c). Macrophages are known to accumulate in adipose tissue in obesity [30] and RNAseq analysis showed reduced expression of several macrophage markers (*Cx3cr1*, *Trem2*, *Gpnmb*, *Spp1*, *Ccl2*) in the adipose tissue of *ANOV* mice relative to *WT* mice on HFD (Fig. 4b). Therefore, we evaluated H&E-stained sections from epididymal (eWAT) and inguinal (iWAT) white adipose tissue depots. In male *WT* mice, HFD-feeding increased the presence of crown-like structures (Fig. 5a), which are typically characterized as dying adipocytes that are surrounded by macrophages[30]. In contrast, *ANOV* mice seemed to have fewer crown-like structures. Consistent with this, the expression of several macrophage markers (*Adgre1*, *Cd68*, *Cd11c*), the chemokine *Ccl2*, and the pro-inflammatory cytokine TNFα (*Tnf*) was diminished in both the eWAT and iWAT of *ANOV* mice compared to *WT* mice in the HFD group (Fig. 5b and Supplemental Fig. 4). On either diet, the expression of the marker *Cd163,* which has been associated with macrophages displaying an anti-inflammatory phenotype[33], was higher in *ANOV* mice compared to *WT* mice (Fig. 5b). Overall, these data indicate that NAMPT overexpression reduces the inflammatory phenotype in eWAT that is induced by diet-induced obesity.

**Fig. 5.**
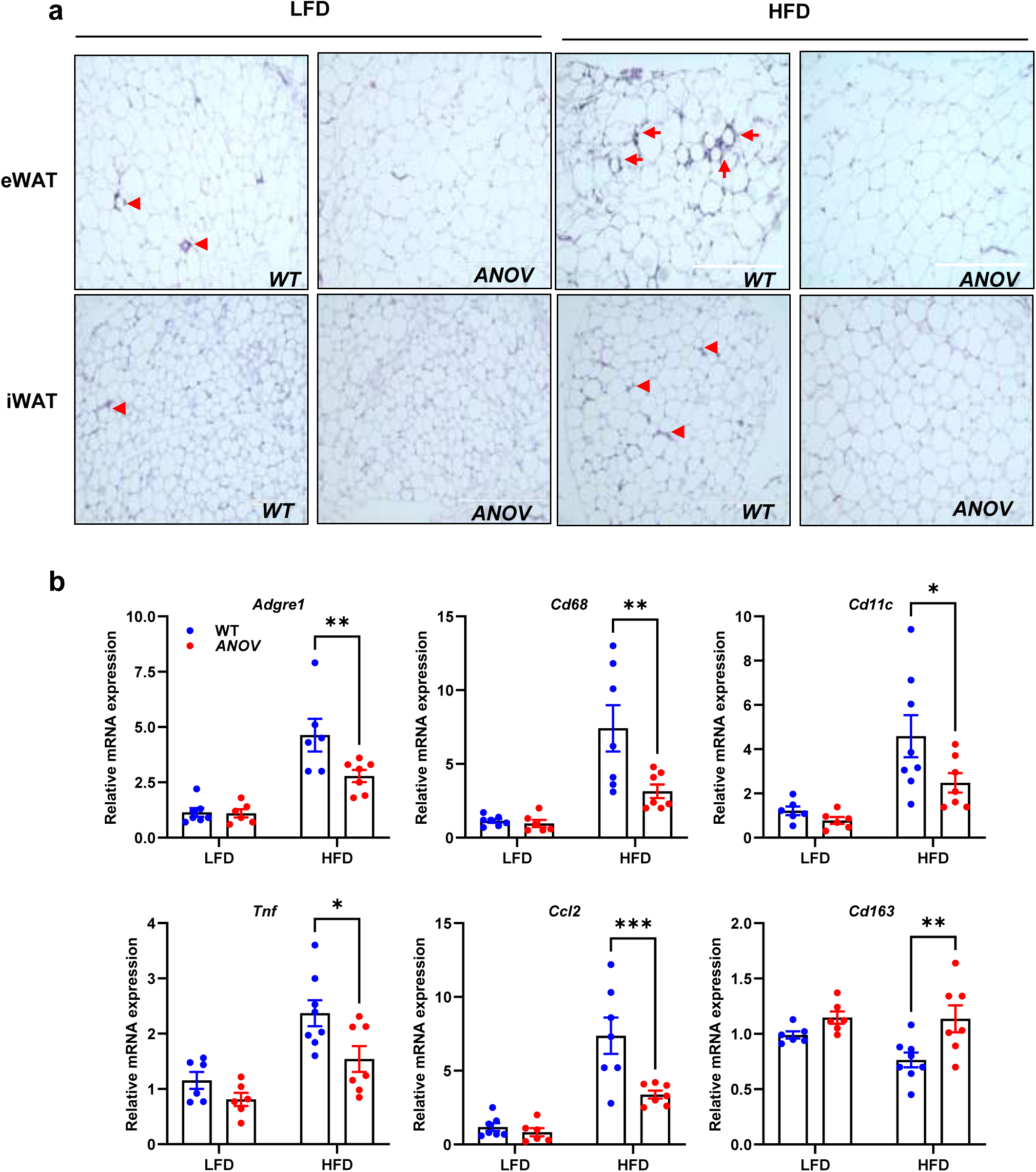
Adipose tissue inflammation is diminished in *ANOV* mice on a high fat diet. **(a).** Representative histological images of eWAT and iWAT after 12 weeks of diet feeding. Arrows indicate crown-like structures. Scale bar = 400 µm. **(b)**. Relative mRNA expression of macrophage markers and pro-inflammatory cytokines were determined in eWAT *WT* and *ANOV* mice by quantitative RT-PCR (LFD n=6-7; HFD n=7-8). The number of mice (n) is presented as individual datapoints. Mean ± s.e.m. shown within dot plots. For multiple comparisons, two-way ANOVA with Holm-Sidák multiple comparison test was performed.*P < 0.05, **P < 0.01, ***P < 0.001.

### ANOV mice are protected from hepatic steatosis and injury due to high-fat diet feeding

Diet-induced obesity is often associated with hepatic metabolic dysfunction and metabolic dysfunction-associated steatotic liver disease (MASLD). Male *ANOV* mice had reduced liver to body weight ratios compared to *WT* mice on either diet (Fig. 6a). In the HFD group, hepatic triglyceride content decreased by 25% in *ANOV* mice (Fig. 6b), and H&E-stained liver sections revealed the appearance of fewer lipid droplets in *ANOV* mice relative to *WT* mice (Fig. 6c). Additionally, circulating alanine transaminase levels, which are a surrogate marker of liver injury, were 85% lower in *ANOV* mice than *WT* in the HFD group (Fig. 6d). Expression of markers associated with stellate cell activation (*Serpine1* and *Acta2*), decreased in HFD-fed *ANOV* mice compared to *WT* mice (Fig. 6e), whereas several genes encoding factors associated with mitochondrial metabolism (*Ppargc1a*, *Cpt1a*, *Ndufb10*, and *Cox7a1*) were increased (Fig. 6f).

**Fig. 6.**
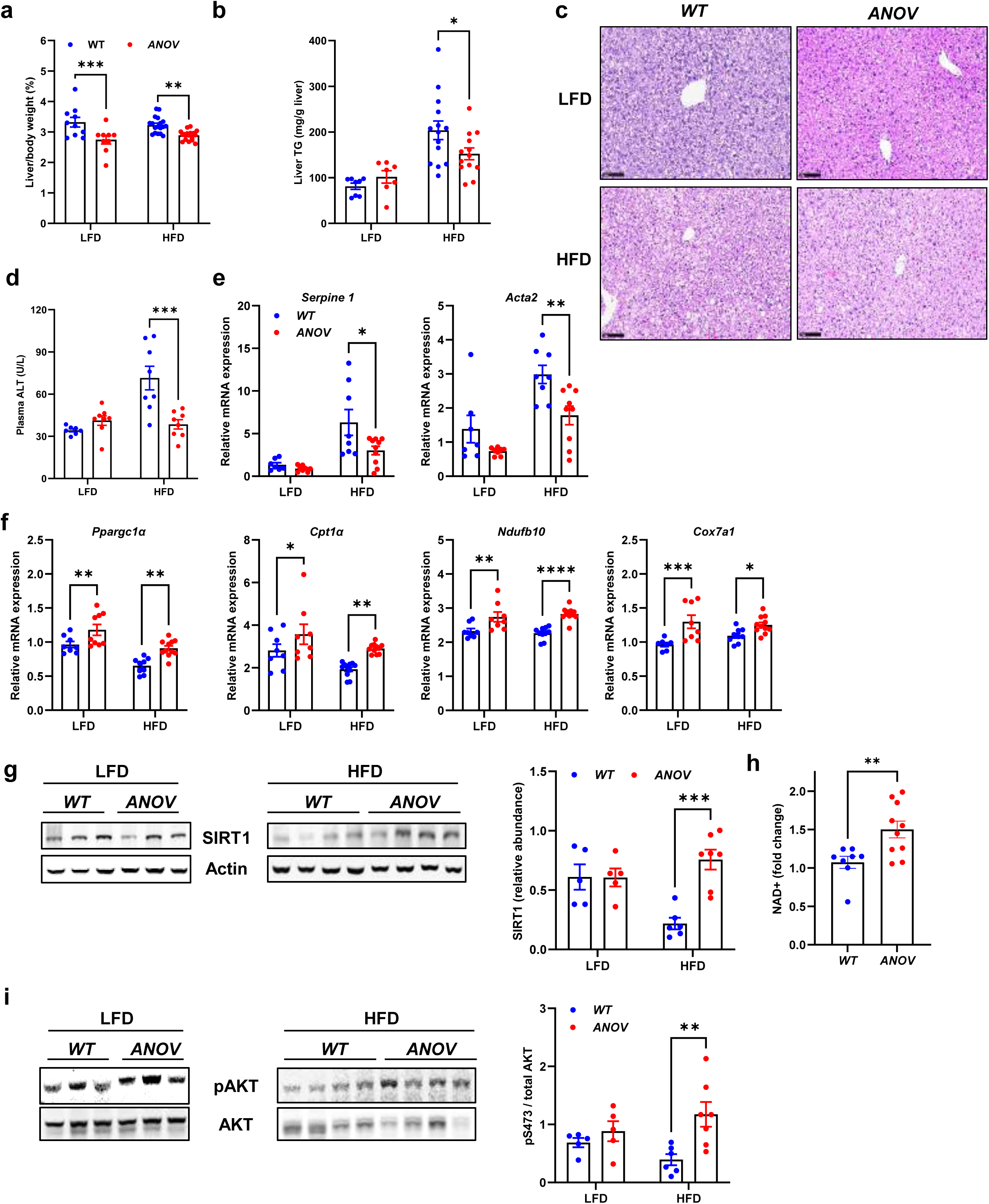
*ANOV* mice are protected from hepatic damage due to high-fat diet feeding. *WT* and *ANOV* male mice were fed with LFD and HFD for 12 weeks followed by subsequent analyses. **(a-b)** Liver size expressed as percent body weight and liver triglycerides (TG) of diet-fed mice (LFD n=7-9; HFD n=14-18). **(c)**. H&E stained liver sections after diet treatment. Scale bar = 100 µm. **(d)**. Plasma ALT levels were measured enzymatically. **(e-f).** Hepatic mRNA expression of genes involved in liver fibrosis (*Serpine1 and Acta2*) and mitochondrial metabolism (*Ppargc1a, Cpt1a, Ndufb10, Cox7a1*) determined by quantitative RT-PCR (LFD n=7-9; HFD n=8-11). **(g).** Immunoblot of liver tissue showing SIRT1 protein levels. Actin was used as a loading control. The graph depicts relative quantification of SIRT1. **(h).** Liver NAD+ levels were determined in HFD fed *WT* and *ANOV* male mice (n=8-10). **(i).** Immunoblots showing phospho-(S473) and total AKT levels in liver of 12-week diet fed *WT* and *ANOV* male mice (LFD n=5; HFD n=6-7). The graph depicts the relative quantification of p/T-AKT. The number of mice (n) is presented as individual datapoints. Mean ± s.e.m. shown within dot plots. For multiple comparisons, two-way ANOVA with Holm-Sidák multiple comparison test was performed. *P < 0.05, **P < 0.01, ***P < 0.001, ****P < 0.0001.

*ANOV* mice exhibited an increased hepatic abundance of SIRT1, a key NAD+-dependent enzyme that controls various metabolic responses and can be positively regulated by NAD+ abundance (Fig. 6g)[34]. Consistent with this, in HFD-fed mice, hepatic NAD+ content was increased in *ANOV* mice compared to controls (Fig. 6h). We also examined hepatic insulin signaling by measuring the phosphorylation of AKT at serine 473. AKT phosphorylation was increased in the liver of *ANOV* mice compared to *WT* mice on HFD (Fig. 6i). In contrast, phosphorylation of AKT in the eWAT or muscle in male *WT* or *ANOV* mice was not altered (Supplementary Fig. 5 a-b). Collectively, these data suggest that adipocyte-specific NAMPT overexpression protects against metabolic liver disease and enhances hepatic NAD+ availability in male mice.

### Adipocyte NAMPT regulates systemic metabolism by EV release

NAMPT overexpression in adipocytes improved metabolic parameters, insulin signaling, and NAD+ content in the liver, possibly suggesting interorgan crosstalk through secretion of NAMPT from adipocytes in extracellular vesicles (EV) [3, 4, 17]. We therefore characterized EV from plasma of *WT* and *ANOV* male mice. Plasma EV had an average diameter of about 100 nm and contained the common EV (exosome) markers ALIX and TSG101, but not the nuclear marker Lamin (Supplemental Fig. 6a). We then determined the total number of EVs and percentage of EV containing NAMPT in plasma by FACS analysis (Supplemental Fig. 6b). The number of plasma EV particles was significantly reduced in male *ANOV* mice compared to *WT* controls (Fig. 7a). However, there was a 3.8-fold increase in the percentage of EV that contained NAMPT in male HFD fed *ANOV* mice (compared to *WT* HFD mice) (Fig. 7b). On either LFD or HFD, female *ANOV* and *WT* mice had a statistically similar number of plasma EV particles (Fig. 7a). The percentage of plasma EV containing NAMPT was higher in female compared to male mice overall (Fig. 7b) and no differences between either genotype was detected for the percentage of EV containing NAMPT in female mice (Fig. 7b). Next, we collected EV released by adipose tissue explants from male *WT* and *ANOV* mice maintained on a HFD for 12 weeks and administered an equal number of EV to HepG2 hepatoma cells. We found that EV from *ANOV* mice enhanced insulin-stimulated AKT phosphorylation signaling in HepG2 cells compared to EV from *WT* mice on HFD (Fig. 7c).

**Fig. 7.**
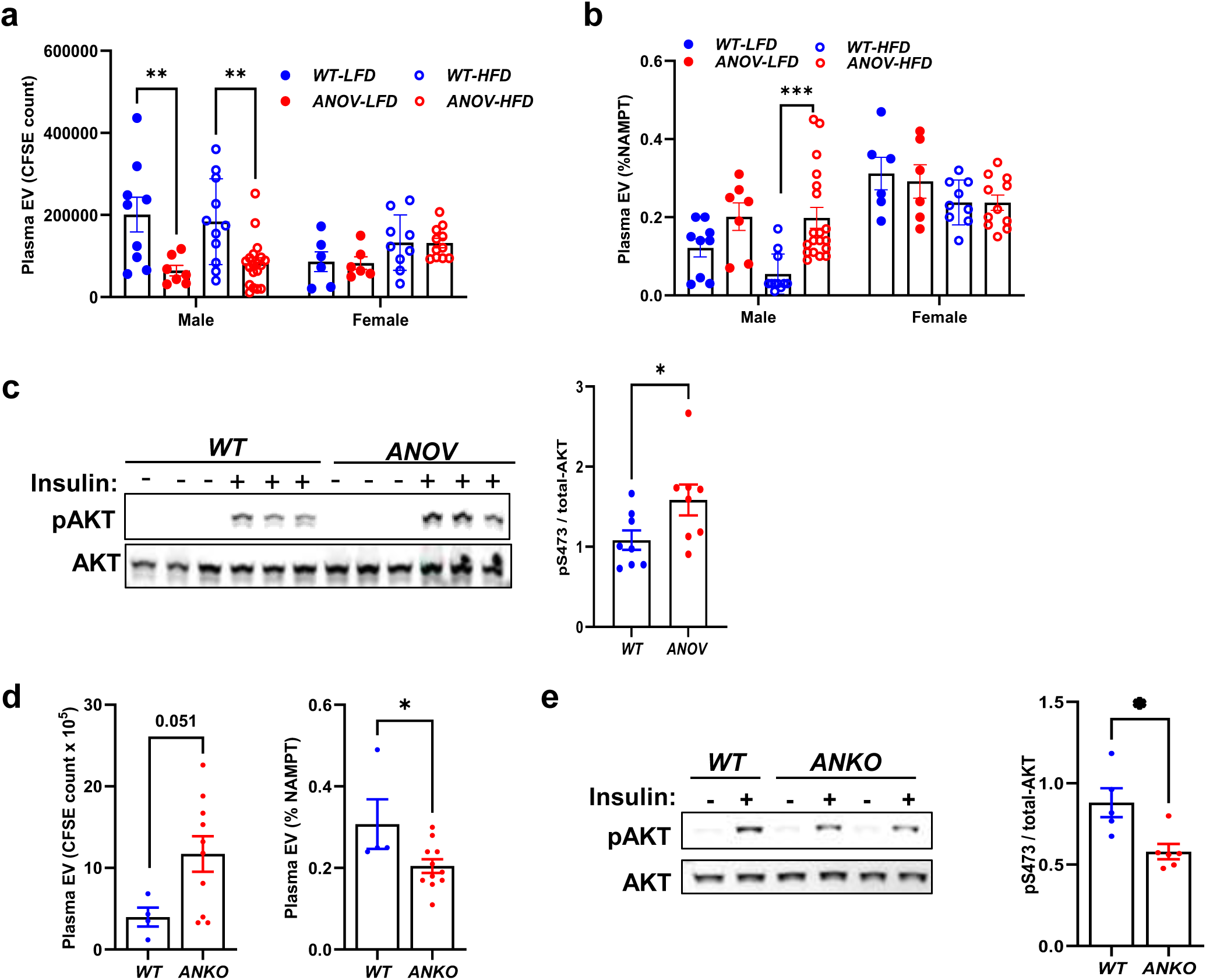
Adipocyte NAMPT regulates insulin signaling by EV release. **(a-b)** Plasma of LFD or HFD fed male and female *WT* or *ANOV* mice was subjected to gel filtration chromatography followed by carboxyfluorescein diacetate succinimidyl ester (CFSE) staining to determine the number of extracellular vesicles (EV) and percent of NAMPT-containing EVs (LFD-male 7-9, female n=6; HFD-male n=11-19, female 9-11).**(c).** EV were isolated from adipose tissue (AT) explants of HFD fed male *WT* or *ANOV* mice and used to treat hepatocytes (HepG2). After 48 hours, cells were serum-starved for 4 hours, then stimulated with insulin (100 nM) for 5 minutes and subjected to immunoblotting for phospho- (S473) and total AKT levels. The graph depicts relative quantification of p/T-AKT. **(d).** Nano FACS analysis showing total number of EVs and percent of NAMPT-containing EVs in plasma of chow-fed female *WT* (n=4) and *ANKO* (n=11) mice (4-month old). **(e).** HepG2 cells were treated with EV isolated from adipose tissue (AT) explants of chow-fed female *WT* and *ANKO* mice (4-month old), then stimulated with insulin and subjected to immunoblotting for phospho- (S473) and total AKT levels. The graph depicts relative quantification of p/T-AKT. The number of mice (n) is presented as individual datapoints. Mean ± s.e.m. shown within dot plots. For multiple comparisons, two-way ANOVA with Holm-Sidák multiple comparison and for two independent data sets, Two-tailed unpaired Student’s t-test was performed. *P < 0.05, **P < 0.01, ****P < 0.0001.

We also examined the effects of EV from adipocyte-specific NAMPT knockout (*ANKO*) mice on insulin signaling. Chow fed female *ANKO* mice, which are insulin resistant[4, 13], had increased numbers of circulating and adipose tissue derived EV compared to *WT* mice (Fig. 7d and Supplemental Fig. 6c). This is consistent with prior work showing that metabolic stress increases adipocyte EV release [35]. Compared to *WT* mice, the percentage of NAMPT-containing EV was reduced in the plasma of chow fed female *ANKO* mice (Fig. 7d). When we treated HepG2 cells with EV collected from female mouse gonadal WAT explants, we found that insulin-stimulated AKT phosphorylation was significantly lower in cells treated with EV from *ANKO* explants compared to *WT* EV treatment (Fig. 7e). Collectively, our data suggests that NAMPT released from adipose tissue EV may enhance insulin signaling on peripheral tissues to improve whole body glucose metabolism.

To investigate whether *ANOV* mouse adipose tissue-derived EV could impact metabolic homeostasis *in vivo*, we collected EV from adipose tissue explants isolated from chow fed *WT* and *ANOV* male mice and administered equal numbers of EV to C57BL6/J mice on HFD. Treatment with *ANOV* EV significantly improved glucose tolerance and lowered plasma TAG compared to *WT* EV treated mice (Fig. 8a-b). EV treatment did not affect body weight (data not shown) or liver AKT-phosphorylation (Fig. 8c). However, plasma ALT was significantly reduced (Fig. 8d) and hepatic SIRT1 expression was significantly enhanced (Fig. 8e) by *ANOV* EV treatment (compared to *WT* EV).

**Fig. 8.**
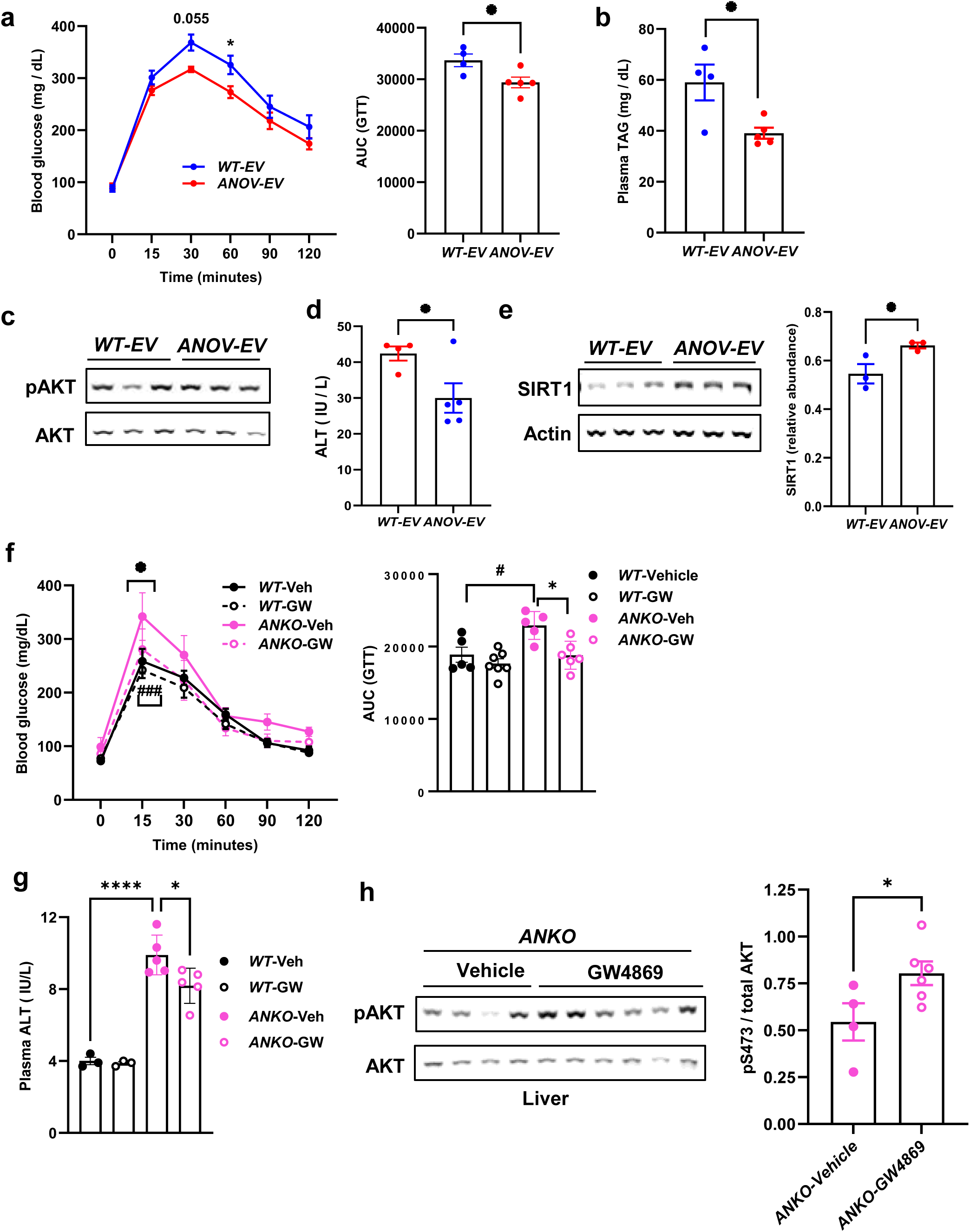
Adipocyte NAMPT containing EV regulates systemic metabolism. **(a-b).** Adipose tissue derived EVs from Chow fed *WT* or *ANOV* mice (4-months old) were intravenously administered to *WT* HFD male mice for 3 weeks followed by performing GTT or plasma TAG (*WT*-EV n=4; *ANOV*-EV n=5), **(c)**. Immunoblot showing phospho- (S473) and total AKT levels in liver of above EV treated mice. The graph depicts relative quantification of p/T-AKT (n=3 / group). **(d)**. Plasma ALT levels were measured in the above mice (n=4-5 / group). **(e)**. SIRT1 expression was determined in liver of above EV treated mice. Actin used as loading control and graph showing relative quantification (n=3 / group). **(f)** *WT* or *ANKO* female mice (4-month old) intraperitoneally injected with Vehicle or GW4869 for 6-weeks followed by performing GTT (*Veh* n=5-7; *GW4869* n= 5-6). **(g)** Plasma ALT was measured in vehicle or GW4869 treated above mice (*Veh* n=3; *GW4869* n= 5). **(h)** Immunoblot showing phospho- (S473) and total AKT levels in liver of above *vehicle* or *GW4869* treated *ANKO* female mice. The graph depicts relative quantification of p/T-AKT (*Veh* n=4; *GW4869* n= 6). The number of mice (n) is presented as individual datapoints. Mean ± s.e.m. shown within dot plots. For multiple comparisons, two-way ANOVA with Holm-Sidák multiple comparison and for two independent data sets, Two-tailed unpaired Student’s t-test was performed. *P < 0.05, ^#^P<0.05, ****P < 0.0001, ^###^P<0.001.

Finally, we treated *ANKO* female mice with GW4869, a compound that suppresses EV release[22]. Sixteen-week-old *WT* and *ANKO* female mice were injected with vehicle or GW4869 intraperitoneally twice a week for 6 weeks. Consistent with prior reports [2–4, 13, 15, 18], *ANKO* vehicle treated mice were glucose intolerant compared to *WT* vehicle controls (Fig. 8f). Glucose tolerance was not affected by GW4869 treatment in *WT* mice. In *ANKO* mice, GW4869 administration improved glucose tolerance compared to vehicle controls (Fig. 8f). GW4869 treatment significantly enhanced hepatic AKT-phosphorylation and decreased plasma ALT in *ANKO* mice (Fig. 8g-h). Collectively, these results suggest that the regulation of adipocyte EV release and contents by NAMPT is crucial for regulating hepatic metabolic function.

## Discussion

Herein, we examined the role of adipocyte NAMPT in regulating systemic metabolic dysfunction in diet-induced obese mice by using novel mice that overexpress NAMPT in WAT. Male *ANOV* mice were protected from diet-induced glucose intolerance and liver metabolic dysfunction independently of changes in body weight or adiposity. However, in female *ANOV* mice, we observed only modest improvements in metabolic phenotypes, likely due to higher NAMPT expression in female mouse adipose tissue. We also determined that a higher percentage of EV from adipose tissue of *ANOV* mice contained NAMPT and that *ANOV*-EV enhanced insulin-stimulated AKT phosphorylation in HepG2 cells and improved glucose tolerance in obese C57BL6/J mice *in vivo*. In contrast, adipose tissue EV from NAMPT-deficient *ANKO* mice impaired insulin signaling *in vitro* and blocking EV release in *ANKO* mice improved glucose tolerance and hepatic AKT phosphorylation *in vivo*. In summary, we identified a novel mechanism by which adipocyte NAMPT activity can regulate systemic metabolic function by modulating the release of EV that can act on peripheral tissues to enhance insulin signaling and improve overall glucose homeostasis and metabolic dysfunction.

Adipose tissue dysfunction can affect whole-body metabolism and is a hallmark of obesity-related disease [36, 37] including insulin resistance and MASLD. Adipocytes release various factors including lipids, hormones, and cytokines that affect systemic metabolism, but recent studies have revealed the importance of adipose-derived EV in metabolic dysfunction[38]. Metabolically-stressed adipocytes exhibit increased release of EV [39], and administration of EV from obese, insulin-resistant mice is sufficient to cause insulin resistance and metabolic dysfunction in lean mice[39, 40]. Similarly, previous studies also demonstrated that plasma EV concentrations were higher in people with obesity and metabolic syndrome [41] and in people with obesity and type 2 diabetes [42] than in healthy lean subjects. Moreover, EV isolated from people with obesity impaired insulin signaling in myotubes in vitro whereas EV from the same subjects after marked weight loss did not[43]. Several microRNAs contained in EV have been implicated in the development of insulin resistance in mouse and human EV[44]. However, a complete understanding of the EV cargoes that are sufficient to confer or prevent metabolic dysfunction remains elusive

Prior work has shown that enzymatically active NAMPT can be released into the bloodstream in EV, internalized into peripheral tissues, and increase NAD+ biosynthesis and SIRT1 expression in recipient cells [17–19, 45]. Consistent with this, we found that male *ANOV* mice had a higher proportion of circulating EV containing NAMPT and had increased SIRT1 protein and NAD+ accumulation in liver. Adipose tissue-derived *ANOV* EV were sufficient to enhance insulin signaling in liver cells and improve glucose tolerance in diet-induced obese mice. In contrast, treatment with adipose tissue-derived EV from *ANKO* mice, which had a lower percentage of NAMPT-containing EV, impaired insulin signaling in HepG2 cells compared to *WT* EV. Furthermore, the inhibition of EV-secretion in *ANKO* mice improved glucose tolerance and AKT-activation in liver. It is tempting to speculate that the transportation of NAMPT itself mediates the sensitizing effects of the EV cargo since SIRT1 is known to enhance insulin signaling in multiple cell types[46, 47], but we cannot exclude the possibility that other *ANOV* EV cargoes mediate these effects. It is also unclear if lower abundance of NAMPT in *ANKO* (compared to *WT*) EV is sufficient to explain diminished insulin signaling in response to *ANKO* EV, or if these EV contain other cargoes that impair the response to insulin. Whether human NAMPT also plays a critical role in regulating EV release and cargo contents and the identification of EV cargos regulated by NAMPT in adipocytes that are sufficient to enhance or impair insulin signaling will require further exploration.

*ANOV* mice exhibited several significant improvements in liver metabolic dysfunction, including reduced hepatic steatosis, lower plasma ALT activity, and enhanced AKT signaling. In contrast, we did not detect enhanced AKT signaling in muscle or adipose tissue of *ANOV* mice. Prior work has shown that liver takes up a high proportion of adipocyte-derived EV[22]. It is possible that the improved insulin signaling in liver, but not muscle or adipose tissue, is due to robust EV uptake by liver compared to other organs or if this is due to tissue-specific effects of the *ANOV* EVs.

We also noted a sex difference in the response to NAMPT overexpression wherein the effects of *ANOV* genotype on glucose and insulin homeostasis were more dramatic in male mice versus female mice. Conversely, prior work has shown that female *ANKO* and germline heterozygous NAMPT KO mice are glucose intolerant compared to *WT* mice, but male mice of either line were not affected[13, 48]. Thus, while the effects of adipocyte NAMPT overexpression are more evident in male mice, the effects of NAMPT deficiency manifest more strongly in female mice. Compared to *WT* male mice, *WT* female mice tend to have increased NAMPT expression in WAT and exhibit much higher circulating NAMPT-containing EV, which could obscure the effects of NAMPT overexpression. Indeed, the transgenic approach to overexpress NAMPT did not result in a significant increase in NAMPT in WAT or circulating EV in female mice likely due to higher endogenous levels. Strikingly, *WT* female mice exhibited a higher percentage of NAMPT-containing EV than male *ANOV* mice (Fig. 7b). It could be speculated that higher circulating NAMPT levels in female mice compared to male mice plays a role in their sex-dependent protection from developing metabolic disease [20, 21]. Consistent with the sex difference in NAMPT expression, the present data also support a role for estrogen in regulating adipose NAMPT expression. Adipose tissue NAMPT protein abundance was diminished after ovariectomy while estradiol supplementation enhanced NAMPT expression in female mice fed HFD (Fig. 3). Estradiol also significantly increased the percentage of plasma EV containing NAMPT (Supplemental Fig. 6d). Altogether, these findings suggest that female sex and estradiol may play an important role in regulating adipose tissue NAMPT expression and release in EV.

In conclusion, we show that overexpression of NAMPT in adipocytes is sufficient to protect male mice from systemic HFD-induced metabolic dysfunction. The beneficial effects include protection from adipose tissue inflammation, hepatic steatosis, glucose intolerance, and impairments in insulin signaling in liver on HFD. Our data suggests that effects of modulating NAMPT in adipocytes are mediated, at least in part, by altering the cargo of adipose tissue-derived EV. Additional studies will be needed to determine whether the release of NAMPT itself mediates these effects, and to understand how this EV cargo alters insulin signaling and metabolic health.

## Acknowledgements

This work was supported by National Institutes of Health (NIH) grants P30 DK056341 (Pilot and Feasibility award to SM), R01 DK104995 to B.N.F., K01 DK137050 (to DF), K08 DK131255 (KL). This research was supported by the Cores of the Washington University Nutrition Obesity Research Center (P30 DK056341), Diabetes Research Center (P30 DK020579), and Digestive Diseases Research Core Center (P30 DK052574). We also thank the Genome Engineering and iPSC Center (GEiC) at the Washington University in St. Louis for gRNA reagent validation services. We thank the Genome Technology Access Center (GTAC) at the McDonnell Genome Institute at Washington University School of Medicine, which is partially supported by NCI Cancer Center Support Grant #P30 CA91842 to the Siteman Cancer Center from the National Center for Research Resources (NCRR), a component of the NIH Roadmap for Medical Research. This publication is solely the responsibility of the authors and does not necessarily represent the official view of NCRR or NIH. This work was also supported by the Hope Center Alafi Neuroimaging Lab at Washington University School of Medicine (Grant No. S10 RR027552) for liver H&E slide imaging.

## Author contributions

D.F., E.G., M.L., M.H., S.T., X.C., E.K., Q.G., E.F., J.K., T.P., K.L., C.C., and S.M.: performed experiments. D.F., X.C., S.T., E.K., Q.G., C.C., and S.M. were responsible for methodology. S.M. were responsible for conceptualization. B.N.F. and S.M., prepared the original draft of the manuscript. D.F., K.M., C.C., J.P.T., G.P., B.N.F., and S.M. reviewed and edited the manuscript. D.F., B.N.F. and S.M., performed the formal analysis. B.N.F., J.Y., and S.M., acquired funding and supervised the study. All authors take full responsibility for all the content of this publication. S.M. is the guarantor of this work and, as such, had full access to all the data in the study and takes responsibility for the integrity of the data and the accuracy of the data analysis.

---

**Supplemental Table 1:**
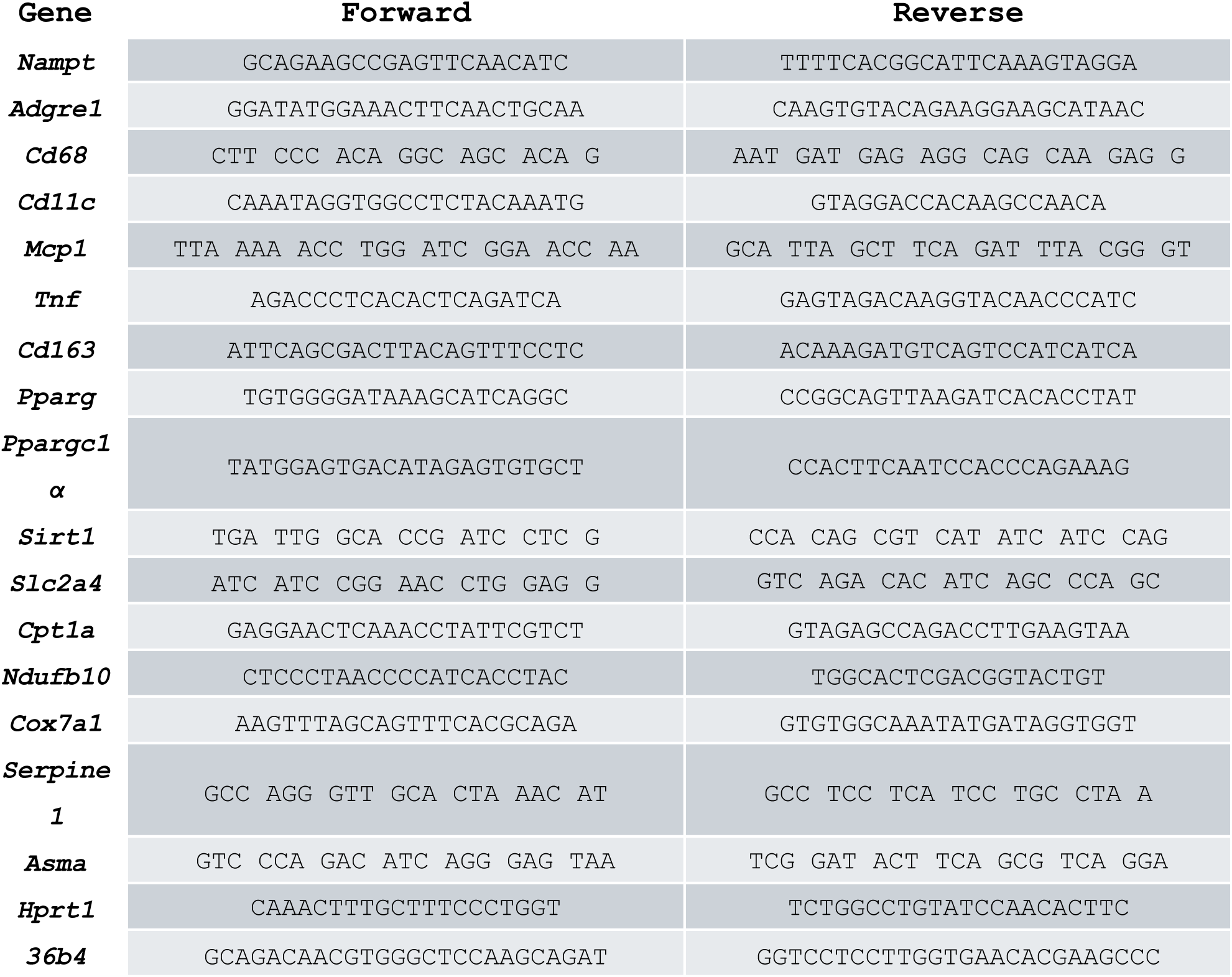
Quantitative RT-PCR primers.

**Supplemental Figure 1.**
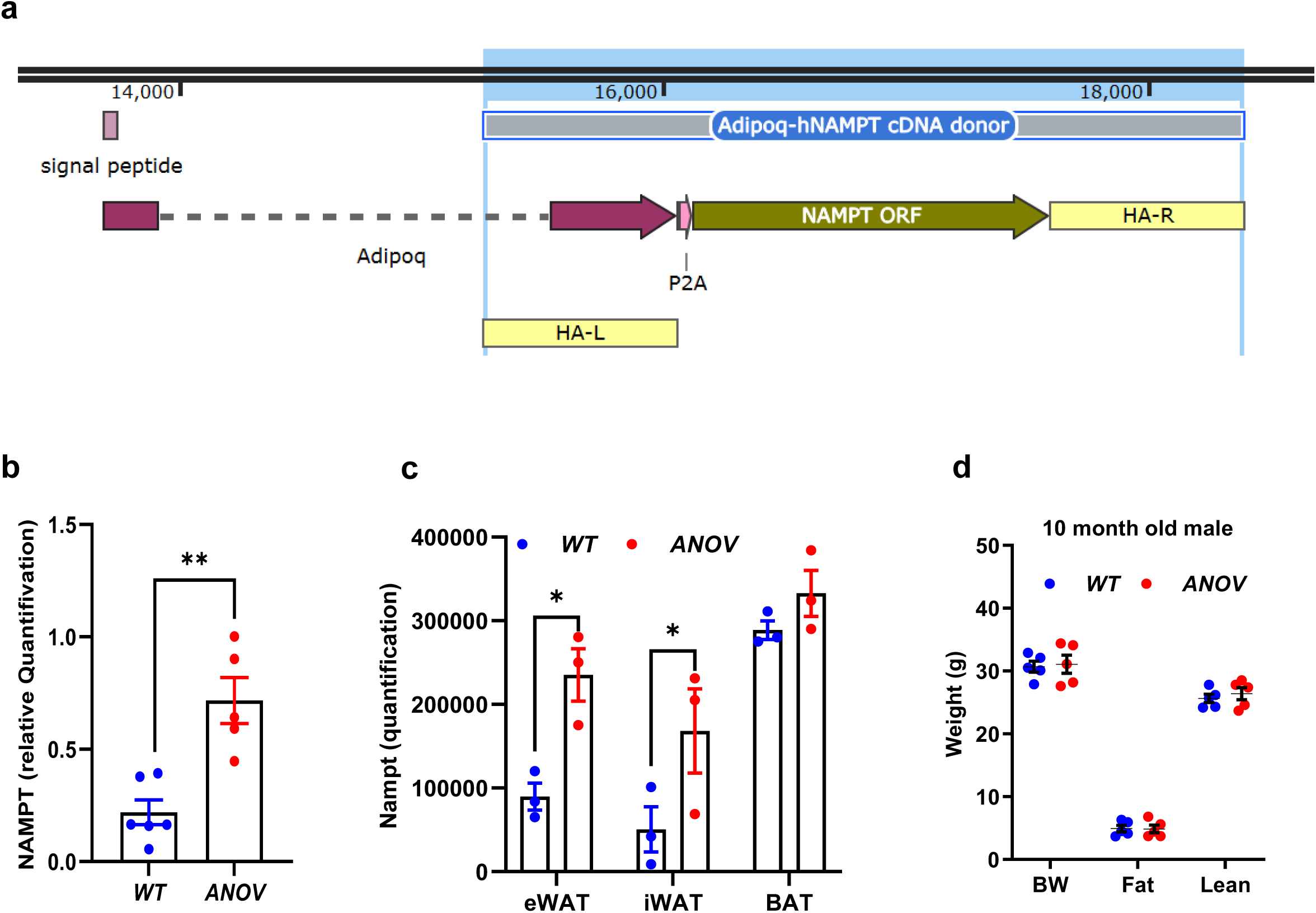

**Supplemental Figure 2.**
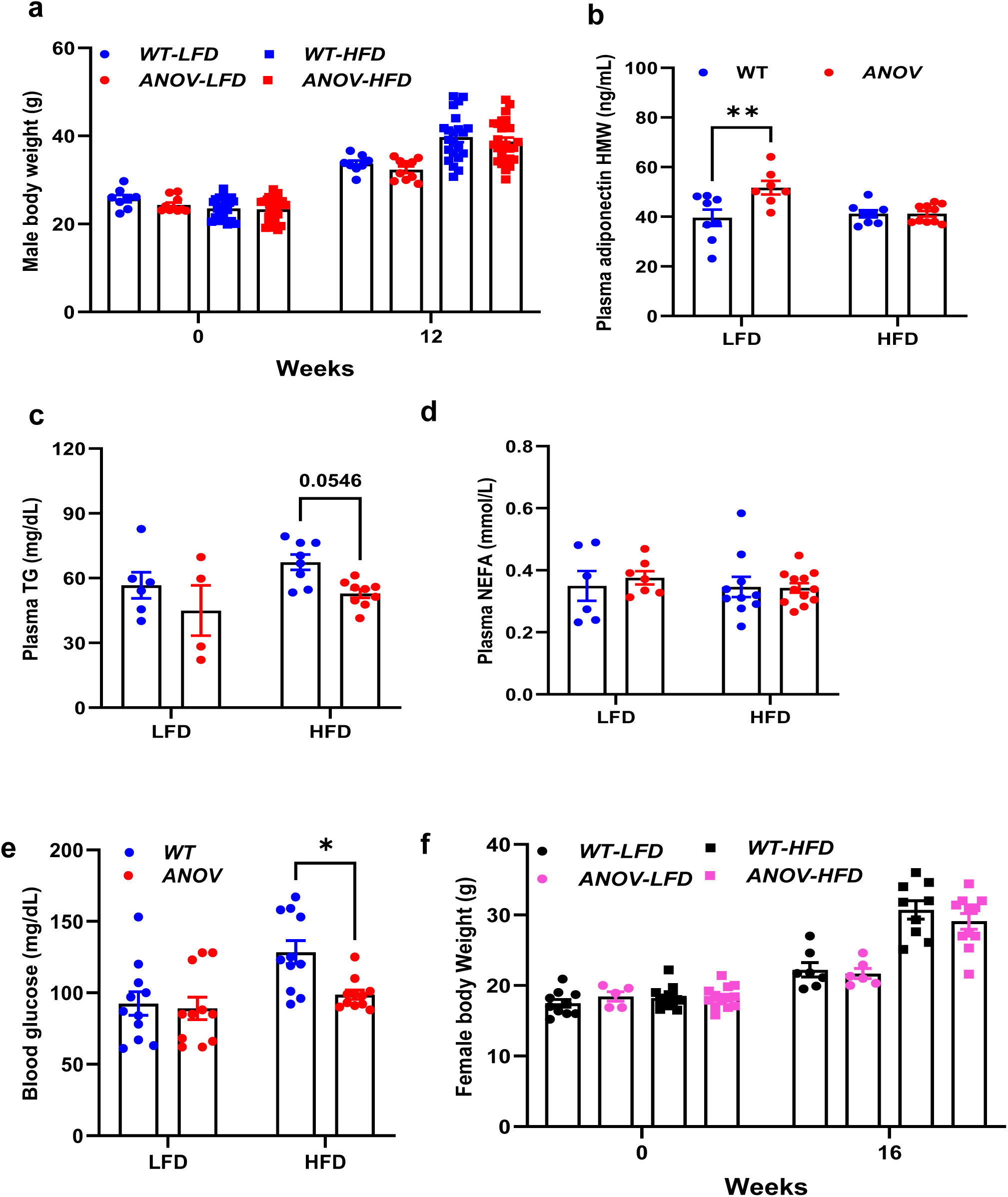

**Supplemental Figure 3.**
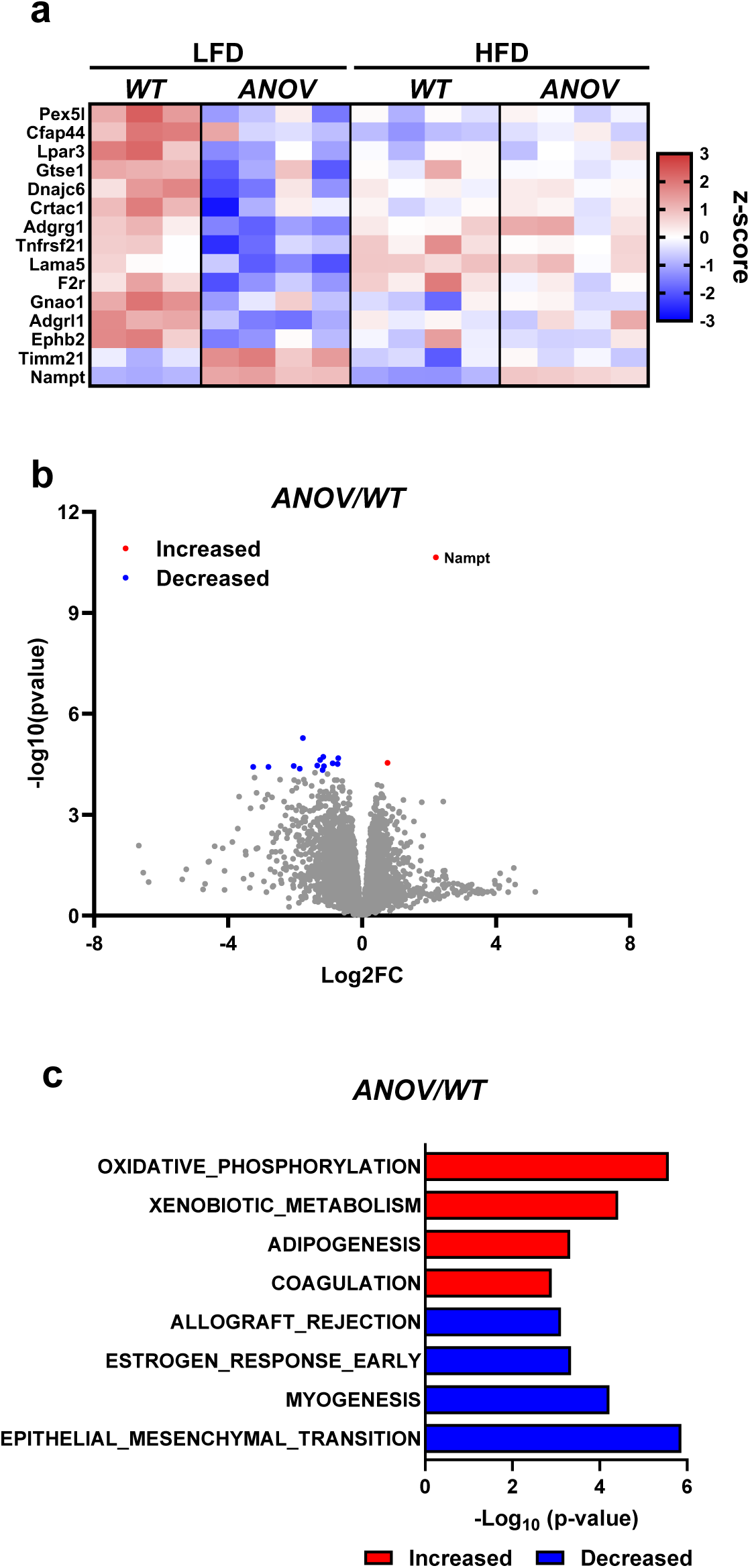

**Supplemental Figure 4.**
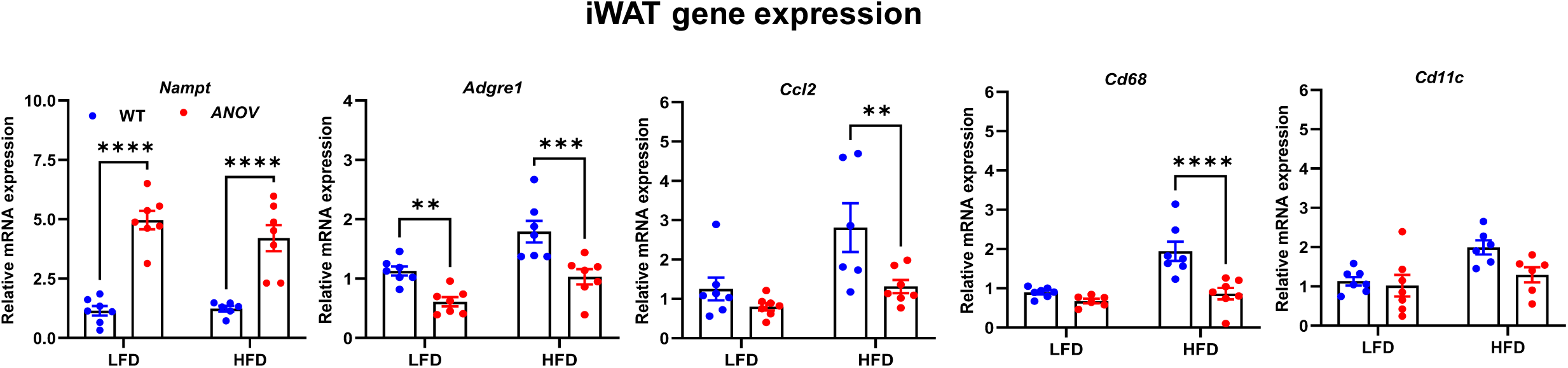

**Supplemental Figure 5.**
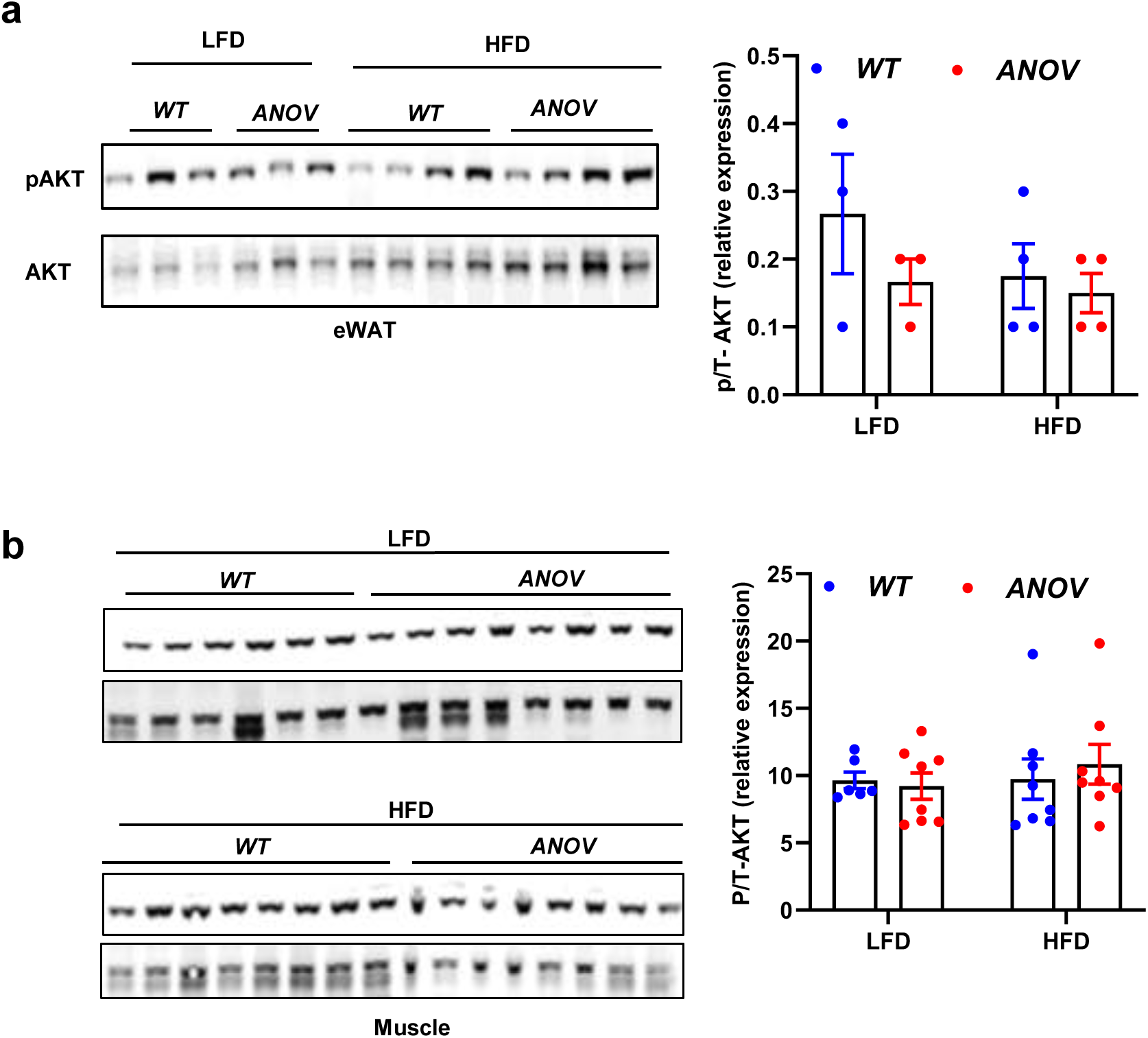

**Supplemental Figure 6.**
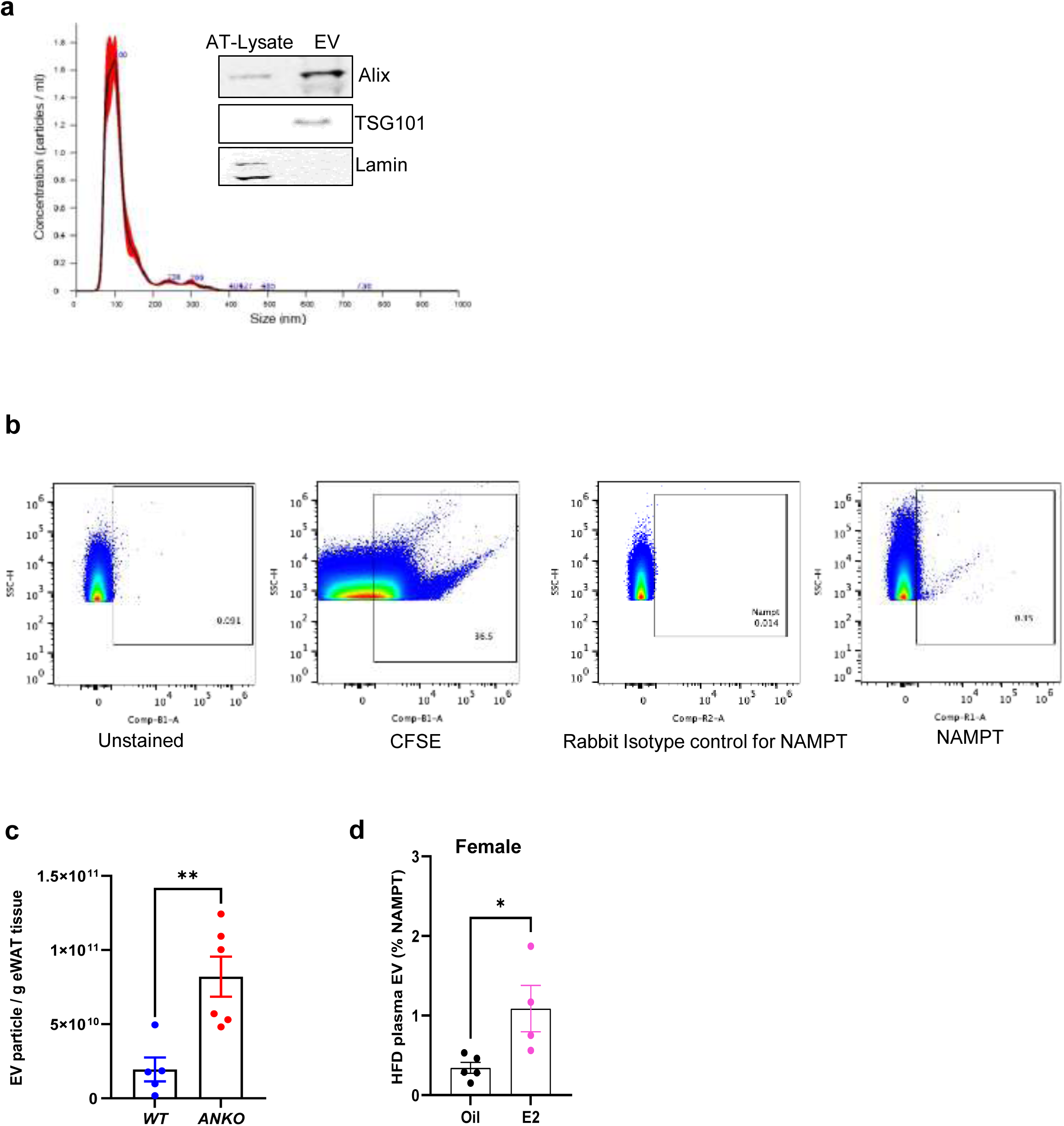

