## Supplementary methods and legends for "Adipose Tissue Overexpression of Nicotinamide Phosphoribosyltransferase Prevents Metabolic Dysfunction in Obese Mice"

**Supplementary legends:**

**Supplemental Fig. 1.** Generation of adipocyte NAMPT overexpressing (*ANOV*) mice.

**(a).** Schematic for generation of adipocyte specific NAMPT overexpressing mice. **(b).** Graph showing relative quantification of eWAT NAMPT vs actin in *WT* and *ANOV* mice. (n=5-6) (related to **Fig 1a**) **(c)**. Graph showing quantification of NAMPT in eWAT, iWAT and BAT fat pads in *WT* and *ANOV* mice. (n=3) (related to **Fig 1b**). **(d)** Body composition of 10-month old male mice on chow diet, determined by ECHO MRI (n=4-5). Data represented as mean ± SEM. Significance determined by Student’s t-test. **P < 0.01.

**Supplemental Fig. 2.** Body weights and plasma parameters in *ANOV* mice on LFD and HFD.

**(a)** Male body weight at week 0 and 12 after diet feeding (LFD n=8-10; HFD n=22-27). **(b-d)** ELISA showing abundance of high molecular weight (HMW) adiponectin in plasma of mice after 12 weeks of diet. Plasma triglyceride and NEFA concentrations determined enzymatically. **(e)** Blood glucose levels were determined in 16-h fasted *WT* or *ANOV* mice (LFD and HFD n=11 mice/genotype). **(f)** Body weight in female mice at week 0 and 16 of diet feeding (LFD n=8-10; HFD n=22-27). The number of mice (n) is presented as individual datapoints. Mean ± s.e.m. shown within dot plots. For multiple comparisons, two-way ANOVA with Holm-Sidák multiple comparison test was performed. *P < 0.05,**P < 0.01, ****P < 0.0001.

**Supplemental Fig. 3.** RNA sequencing reveals increased adipogenesis *ANOV* adipose tissue on LFD.

Bulk RNA sequencing was performed on eWAT from *WT* and *ANOV* male mice after 12 weeks of either LFD or HFD.

**(a)** Heatmap representation of gene expression across all groups of genes that were significantly differentially expressed (adj. P value < 0.05) in LFD-fed *ANOV* vs LFD-fed *WT*. **(b)** Volcano plot of differentially expressed genes in LFD-fed *ANOV* vs LFD-fed *WT* with genes that were significantly downregulated or upregulated highlighted in blue or red, respectively (adj. P value < 0.05). **(c)** Hallmark gene sets that were significantly different (false discovery rate < 0.05) and substantially increased (red; LogFC > 3) or decreased (blue; LogFC < -3) when comparing LFD-fed *ANOV* vs LFD-fed *WT*.

**Supplemental Fig. 4.** iWAT inflammation is diminished in *ANOV* mice on a high fat diet.

Relative mRNA expression of *Nampt* and macrophage markers (*Adgre1, Ccl2, Cd68, Cd11c*) in iWAT of male mice after 12 weeks of diet (n=6-7). The number of mice (n) is presented as individual datapoints. Mean ± s.e.m. shown within dot plots. For multiple comparisons, two-way ANOVA with Holm-Sidák multiple comparison test was performed. **P < 0.01, ***P < 0.001.

**Supplemental Fig. 5.** Determination of NAMPT protein in eWAT and muscle.

**(a-b)** Immunoblot showing NAMPT protein abundance in Epididymal white adipose tissue (eWAT, LFD n=3; HFD n=4) and muscle (LFD n=6-8; HFD n=8) of diet fed *WT* or *ANOV* male mice. Actin was used as loading control.

**Supplemental Fig. 6.** Isolation of plasma EV.

**(a)** EV size was analyzed using NanoSight nanoparticle tracking analyzer and immunoblot was performed with EV markers (ALIX) and TSG101 and nuclear protein marker Lamin. **(b)** Representative Nano-FACS analysis showing gating for CFSE stained EV positive counts in mice plasma (left two graphs) and in rabbit isotype control and NAMPT containing EVs (right two panels). **(c)** Chow fed female *WT* or *ANKO* mice (4-months old) EV particles were collected from adipose tissue explants and counted in nanoparticle tracker (n= 5-6). **(d)** Plasma of vehicle and estradiol supplemented (E2) HFD female mice showing percent of NAMPT-containing EVs (n=5). The number of mice (n) is presented as individual datapoints. Mean ± s.e.m. shown within dot plots. For multiple comparisons, two-way ANOVA with Holm-Sidák multiple comparison test was performed. **P < 0.01, ***P < 0.001. Significance determined by Student’s t-test. **P < 0.01.

**Supplementary Methods**

**Generation of ANOV mice**

ANOV mice were generated to express a human NAMPT cDNA under the endogenous Adipoq promoter by the Genome Engineering & Stem Cell Center (GESC@MGI) and the Mouse Genetics Core at Washington University in St. Louis. Briefly, the human NAMPT coding sequence was inserted at the C-terminus of the adiponectin (Adipoq) open reading frame (ORF). The P2A peptide was inserted between the Adipoq ORF and NAMPT sequence so that adiponectin and human NAMPT are expressed as two separate proteins. The gRNA was designed to cleave close to the Adipoq stop codon with the recognition sequence: 5’-TATGGGTAGTTGCAGTCAGTTGG (with PAM site underlined). Synthetic gRNA was ordered from Integrated DNA Technologies (Coralville, IA) and validated in N2a cells. In short, the sgRNA was complexed with recombinant cas9 protein and nucleofected into mouse N2a cells. Amplicon NGS detected high percentage of indels at the target site, indicating of efficient cleavage activity. The donor template with two 800 bp homology arms flanking the human NAMPT cDNA was synthesized and cloned into an rAAV backbone by BioBasic (Toronto, Canada). rAAV was made by the Hope Center Viral Core at Washington University in St. Louis. The rAAV donor was also validated by co-delivered into N2a cells along with sgRNA/cas9 complex for targeted integration by junction PCRs. Single-cell mouse embryos incubated with the rAAV donor for 5 hours before being electroporated with Cas9 protein/sgRNA complex. The surviving embryos were transferred into pseudo-pregnant females. Live births were screened for positive insertion junctions to identify founders[1] Primer sequences for 5’ junction are provided in supplementary method.

Primer sequences for 5’ junction:

XCC216a.5Jxn.F- 5’-CTG GAA ATC TGC TGT GCT TTG GG-3’ and XCC216a.5Jxn.R2- 5’-GAG TAA CCT TGT AGG AGT CGG TGG -3’

For 3’ junction: XCC216a.3Jxn.F2- 5’-TGC ACA GCT GAA TAT TGA ACT GGA AG-3’ and XCC216a.3Jxn.R- 5’-GCT GTG TGG AAA GTG TCC GGC-3’

**Estrogen study**: To investigate the effect of estrogen on NAMPT protein expression, 14–17 week-old female C57BL6/J mice were randomized to either sham surgery or bilateral ovariectomy (OVX). Following surgery, mice were placed on HFD (D12451; Research Diets) and after 4 weeks on diet mice were euthanized [2]. To determine the effects of estrogen supplementation, female C57BL6/J mice (22-24-week-old) underwent OVX surgery and were placed on HFD for 10 weeks. After 6 weeks of dietary intervention, OVX mice received either 17β-estradiol (E2) replacement (36 µg/mL) or sesame oil (control) via silastic tubing (20 mm tube, 3 mm surgical glue caps) placed between the scapulae blades for the remaining 4 weeks of intervention. Silastic tubing was replaced every 2 weeks to maintain circulating E2 levels. For both studies (ovariectomy and E2 supplementation), all mice were co-housed (n=4 per cage) at thermoneutrality (30°C) on a reverse light cycle (Dark 10:00-22:00) with *ad libitum* access to water and diet.

**Ex vivo adipose tissue incubation and EV isolation**

Isolation of adipose tissue extracellular vesicles (EV) was performed as previously described [3]. Briefly, after 12 weeks of HFD feeding, mice were euthanized and perfused with 6 mL PBS through the left ventricle of the heart. Epididymal white adipose tissue (eWAT) depots (1 g) were harvested and minced in Hanks’ balanced salt solution (HBSS). The remaining steps in the protocol were conducted under sterile conditions in a laminar flow hood. The minced tissue was transferred to a 100 mm cell strainer and rinsed with 40 mL of sterile HBSS to remove debris from damaged cells. The washed tissue pieces were then transferred to a 10 cm cell culture dish containing cell culture media (FluoroBrite DMEM supplemented with 10% exosome-depleted FBS, EXOFBSHI50A1; System Biosciences) and 1× penicillin/ streptomycin). Tissues were incubated for ~16 h in a cell culture incubator (37°C, 5% CO2). Media were recovered and centrifuged at 600 x *g* for 15 min at 4°C to remove any cells. The supernatant was then centrifuged at 1,200 x *g* at 4°C for 20 min to remove cell debris and apoptotic bodies. The resulting supernatant was centrifuged at 10,000 x *g* for 30 min at 4°C to remove large EV. The final supernatant was concentrated to 1 mL using a centrifugal filter with a 100-KD cutoff (Amicon Ultra-15, UFC910024; MilliporeSigma). The full volume (1 mL) was loaded onto a 10 mL gravity flow size exclusion column (89898; Pierce) packed with Sepharose CL-2B particles (CL2B300; MilliporeSigma) as previously described . Sterile PBS (10 mL) was used to elute EV from the column, and 1 mL fractions were collected. Fractions 2–5 were found to have the highest and purest EV yield, so they were combined and concentrated again to ~300 µL. sEV concentration was determined using the nanoparticle tracking analyzer (NanoSight NS300, Malvern Pananalytical). For western blot and marker analysis, 100 µl of EV suspension was mixed with 10X RIPA buffer and further processed for immunoblotting. EV were isolated and subjected to analysis as per the guidelines of the International Society for Extracellular Vesicles, ISEV [3].

**Glucose and insulin tolerance tests**

Lean mass and fat mass were determined using EchoMRI before tolerance testing to determine lean mass for injections. Before all tolerance tests, mice were placed on hardwood bedding, and glucose tolerance tests were performed in 16-hour-fasted mice. Glucose was dissolved in saline and given via an intraperitoneal injection (2 g glucose/kg lean mass). Insulin tolerance tests were performed in 4-hour-fasted mice. Recombinant human insulin (Humalin R, Eli Lilly; 0.75 U/kg lean mass) in saline was given via an intraperitoneal injection. Blood was procured from the tail vein, and glucose was monitored via a glucometer (Contour Next, Bayer) at the times indicated.

**Liver triglyceride and plasma analyses**

Liver lipid extraction was performed based on previously published paper [4]. Briefly, snap-frozen liver samples were weighed and homogenized in ten volume of ice-cold PBS. Two-hundred microliters of the homogenate was transferred into 1.2 mL of chloroform: methanol (2:1; v/v) mixture followed by vigorous vortexing for 30 s. One-hundred microliters of ice-cold PBS was then added into the mixture and mixed vigorously for 15 s. The mixture was then centrifuged at 4,200 rpm for 10 min at 4°C. Two-hundred microliters of the organic phase (bottom layer) was transferred into a new tube and dried by evaporation. Two-hundred microliters of 1% Triton X-100 in ethanol was used to dissolve the dried lipid with constant rotation for 2 h and TAG was measured.

Blood from 4-hour-fasted mice was collected into EDTA-coated tubes. Plasma was separated via centrifugation at 3,000 X *g* for 10 minutes at 4°C. Plasma and liver triglycerides (TAG) were measured by using commercially available colorimetric assays kit (Thermo Fisher Scientific, TR22421). Plasma non-esterified free fatty acids (NEFAs) were measured using commercial kit (Wako, 999-34691, 995-34791, 991-34891, and 993-35191) as per the manufacturer’s instruction. Plasma alanine transaminase (ALT) and aspartate aminotransferase (AST) were measured using liquid kinetic assays (TECO Diagnostic, A534 and A559). Plasma insulin was determined by Single Plex Immunoassay at the Washington University Core Laboratory for Clinical Studies. Plasma high molecular weight adiponectin was determined using a mouse high molecular weight adiponectin ELISA Kit (Cat # MBS742124-mybiosource).

**Histology**

Tissues harvested as described above were fixed in 10% neutral-buffered formalin for 48 hours. Samples were rinsed and stored in 70% EtOH until paraffin embedding, sectioning, and H&E staining by the Anatomic Molecular Pathology Core Labs at Washington University School of Medicine. Stained sections were imaged with an EVOS FL Color Imaging System (Life Technologies) using the 10X objective.

**Immunoblotting**

Twenty micrograms of protein was loaded onto 4%–15% acrylamide precast gels (Bio-Rad, 64329760) in Laemmli loading buffer (Bio-Rad, 161-0737). Proteins were transferred to PVDF membranes in Tris-glycine buffer with 10% methanol. After transfer, membranes were blocked in 5% BSA in TBS for 1 hour before overnight antibody incubations in 5% BSA in TBS. Antibodies used were as follows: NAMPT (rabbit, Bethyl Laboratories-A300-372A), phospho-AKT-Ser473 (rabbit, CST-4060), AKT (mouse, CST- 2920), ALIX (rabbit, CST-92880), Lamin (rabbit, CST-2032), SIRT1 (rabbit, CST-8469) and β-actin (mouse, CST-3700). After primary antibody incubations, membranes were washed and incubated in secondary antibodies (LI-COR) before imaging on a LI-COR Odyssey.

**mRNA isolation and quantification**

Total RNA was isolated from frozen tissue using Qiazol lysis reagent plus RNeasy Lipid Tissue RNA purification Mini kit (Qiagen, Ref# 1023539) and TRIzol Plus RNA purification kits (Thermo Fisher Scientific, 12183555), according to the manufacturer’s instructions. One µg of RNA was reverse-transcribed into cDNA using High-Capacity reverse transcription kit (Applied Biosystems, 2983336). Quantitative PCR was performed using Power SYBR Green (Applied Biosystems, 4367659) and measured using an ABI QuantStudio 3 sequence detection system (Applied Biosystems). Results were quantified using 2–ΔΔCt and are shown as relative expression with respect to the control groups. Primer sequences are listed in Supplemental Table 1.

**Bulk RNA sequencing**

Bulk RNA sequencing was performed at the Genomic Technologies and Access Center at the McDonald Genomic Institute of Washington University School of Medicine as previously described [5, 6]. Library preparation was performed on 500-1000 ng of total RNA following removal of ribosomal RNA by an RNase-H method using RiboErase kits (Kapa Biosystems). cDNA was synthesized using the SuperScript III RT enzyme (Life Technologies, per manufacturer's instructions). cDNA was blunt ended, had an A base added to the 3' ends, and then had Illumina sequencing adapters ligated to the ends. Sequencing was performed on an Illumina NovaSeq X Plus using paired end reads extending 150 bases. RNA-seq reads were aligned to the Ensembl release 101 primary assembly with STAR version 2.7.9a1 and gene counts were derived from the number of uniquely aligned unambiguous reads by Subread:featureCount version 2.0.32. Total RNA integrity was determined using Agilent Bioanalyzer or 4200 Tapestation. Library preparation was performed with 500ng to 1ug of total RNA. Ribosomal RNA was removed by an RNase-H method using RiboErase kits (Kapa Biosystems). mRNA was fragmented in reverse transcriptase buffer and heating to 94 degrees for 8 minutes. mRNA was reverse transcribed to yield cDNA using SuperScript III RT enzyme (Life Technologies, per manufacturer's instructions) and random hexamers. A second strand reaction was performed to yield ds-cDNA. cDNA was blunt ended, had an A base added to the 3' ends, and then had Illumina sequencing adapters ligated to the ends. Ligated fragments were then amplified for 12-15 cycles using primers incorporating unique dual index tags. Fragments were sequenced on an Illumina NovaSeq X Plus using paired end reads extending 150 bases. Basecalls and demultiplexing were performed with Illumina’s bcl2fastq software with a maximum of one mismatch in the indexing read. RNA-seq reads were then aligned to the Ensembl release 101 primary assembly with STAR version 2.7.9a1. Gene counts were derived from the number of uniquely aligned unambiguous reads by Subread:featureCount version 2.0.32. Isoform expression of known Ensembl transcripts were quantified with Salmon version 1.5.23. Sequencing performance was assessed for the total number of aligned reads, total number of uniquely aligned reads, and features detected. The ribosomal fraction, known junction saturation, and read distribution over known gene models were quantified with RSeQC version 4.04. All gene counts were imported into the R/Bioconductor package EdgeR [7] and TMM normalization size factors were calculated to adjust for samples for differences in library size. Ribosomal genes and genes not expressed in the smallest group size minus one samples greater than one count-per-million were excluded from further analysis. The TMM size factors and the matrix of counts were then imported into the R/Bioconductor package Limma [8]. The performance of all genes was assessed with plots of the residual standard deviation of every gene to their average log-count with a robustly fitted trend line of the residuals. Differential expression analysis was then performed to analyze for differences between conditions and the results were filtered for only those genes with Benjamini-Hochberg false-discovery rate adjusted p-values less than or equal to 0.05. For each contrast extracted with Limma, global perturbations in known MSigDb and KEGG pathways were detected using the R/Bioconductor package GAGE [9] to test for changes in expression of the reported Log2 fold-changes reported by Limma in each term versus the background Log2 fold-changes of all genes found outside the respective term. RNA sequencing data were deposited in the GEO database (GSE239490).

**NAD(H) extraction and quantification**

For mouse tissues, 40 µl of 2:2:1 (v/v/v) methanol:acetonitrile:water with 0.1 M formic acid (pre-cooled at 4 °C) was added per mg of tissue weight. The mixture was homogenized using an Omni Bead Mill homogenizer (OMNI International, Waterbury, CT, USA) at 2.50 m/s for 30 s. 2 M ammonium bicarbonate was added to the samples immediately to adjust the pH to 8.5. The samples were subsequently stored at -20 °C for 1 h. After protein precipitation, samples were centrifuged at 14,000 g at 4 °C for 15 min. The supernatant was transferred to LC-MS vials for same-day LC-MS analysis.

An Agilent quadruple time-of-flight mass spectrometer (6545 Q-TOF) coupled to an Agilent 1290 UHPLC systems was utilized for LC-MS analysis. Metabolites were separated by using a hydrophilic interaction liquid chromatography column (100 x 2.1 mm, 5 μm, 200 Å, iHILIC-(P)Classic with a guard column 20 × 2.1 mm, 5 μm, 200 Å, iHILIC-(P)Classic, HILICON, Sweden). The column compartment temperature was set to 40 °C and the injection volume was 4 μL. Mobile phase A was 95% water and 5% acetonitrile with 20 mM ammonium bicarbonate, 0.1% ammonium hydroxide solution (25% in water), and 2.5 μM medronic acid. Mobile phase B was 95% acetonitrile and 5% water. Separation was carried out using a flow rate of 0.25 mL/min and the following linear gradient: 90% B from 0 to 1 min, 90% to 25% B from 1 to 14 min, 25% B from 14 min to 14.5 min, and 25% to 90% B from 14.5 to 15 min. The column was equilibrated with 90% B for 5 min at a flow rate of 0.4 mL/min followed by 2 min at 0.25 mL/min. MS analysis was operated in negative mode using electrospray ionization. Instrument parameters were as follows: gas, 200 °C at 10 L/min; nebulizer, 44 psi; sheath gas, 300 °C at 11 L/min; capillary, 3000 V; fragmentor, 100 V; skimmer, 65 V; and scan rate, 3 scans per second. The mass range was set to 50 to 1,500 *m*/*z*. Retention time of the authentic standards and accurate mass were used for identification. [M-H]*^-^* ions were used for NAD*^+^* (m/z = 662.1018) and NADH (m/z = 664.1175) to obtain peak areas, and NAD(H) concentrations were determined by an external calibration curve.
